# RyR2 Binding of an Antiarrhythmic Cyclic Depsipeptide Mapped Using Confocal Fluorescence Lifetime Detection of FRET

**DOI:** 10.1101/2023.06.22.546083

**Authors:** Jaroslava Seflova, Jacob A. Schwarz, Abigail N. Smith, Bengt Svensson, Daniel J. Blackwell, Taylor A. Phillips, Roman Nikolaienko, Elisa Bovo, Robyn T. Rebbeck, Aleksey V. Zima, David D. Thomas, Filip Van Petegem, Björn C. Knollmann, Jeffrey N. Johnston, Seth L. Robia, Răzvan L. Cornea

## Abstract

Hyperactivity of cardiac sarcoplasmic reticulum (SR) ryanodine receptor (RyR2) Ca^2+^-release channels contributes to heart failure and arrhythmias. Reducing RyR2 activity, particularly during cardiac relaxation (diastole), is a desirable therapeutic goal. We previously reported that the unnatural enantiomer (*ent*) of an insect-RyR activator, verticilide, inhibits porcine and mouse RyR2 at diastolic (nanomolar) Ca^2+^ and has in vivo efficacy against atrial and ventricular arrhythmia. To determine the *ent*-verticilide structural mode of action on RyR2 and guide its further development via medicinal chemistry structure-activity relationship studies, here we used fluorescence lifetime (FLT)-measurements of Förster resonance energy transfer (FRET) in HEK293 cells expressing human RyR2. For these studies, we used an RyR-specific FRET molecular-toolkit and computational methods for trilateration (i.e., using distances to locate a point of interest). Multi-exponential analysis of FLT-FRET measurements between four donor-labeled FKBP12.6 variants and acceptor-labeled *ent*-verticilide, yielded distance relationships placing the acceptor probe at two candidate loci within the RyR2 cryo-EM map. One locus is within the Ry12 domain (at the corner periphery of the RyR2 tetrameric complex). The other locus is sandwiched at the interface between helical domain 1 and the SPRY3 domain. These findings document RyR2-target engagement by *ent*-verticilide, reveal new insight into the mechanism of action of this new class of RyR2-targeting drug candidate, and can serve as input in future computational determinations of the *ent*-verticilide binding site on RyR2 that will inform structure-activity studies for lead optimization.

## Introduction

The leaky state of the cardiac ryanodine receptor (RyR2 isoform) Ca^2+^-release channels has emerged as a high-potential target for antiarrhythmic drug discovery and development (2). In a healthy heart, cytoplasmic Ca^2+^ during the relaxation phase of the cardiac cycle (diastole) is maintained at nanomolar (nM) concentrations, and Ca^2+^ escape from the sarcoplasmic reticulum (SR) is insignificant. Then, during cardiac contraction (systole), activation of RyR2 results in a massive but transient release of Ca^2+^ from the SR, increasing cytosolic [Ca^2+^] to micromolar (μM) levels. An inappropriate basal leak of Ca^2+^ from the SR via resting RyR2 during diastole is dangerous, as it can lead to plasma membrane depolarization and cardiac arrhythmias (3). Thus, inhibiting the resting RyR2 leaky state could be useful as an anti-arrhythmic therapy (4,5). However, decreasing RyR2 Ca^2+^ flux during systole would have an undesirable side effect of reducing the strength of cardiac contraction. Thus, an ideal antiarrhythmic drug would prevent RyR2 leak during diastole (at nM Ca^2+^), without a negative effect on normal RyR2 Ca^2+^ flux during systole (at μM Ca^2+^).

Drug-like small-molecule modulators of the RyRs have been identified by serendipity, and some of these have been under consideration as inhibitors of the pathologic Ca^2+^-leak through RyR2, with potential uses as antiarrhythmics and in other clinical indications linked to hyperactive resting RyRs (4,5). In a similar manner, we have identified the unnatural enantiomer (*ent*) of verticilide (a natural cyclic depsipeptide produced by the fungus Verticillium sp. (6)), as a specific inhibitor of the RyR2 pathologic leak in nanomolar (i.e., resting) Ca^2+^ (7) and demonstrated efficacy against atrial and ventricular arrhythmias in vivo (7,8). Locating the RyR binding site of such drug candidates, represents a major prerequisite for (a) understanding their mechanism of action, and (b) their further optimization via structure-guided medicinal chemistry.

A major hurdle in locating a small molecule bound to an RyR is the size of the RyR complex. RyRs are large homo-tetrameric channel complexes (∼5000 amino acid residues per protomer) comprised of a small trans-membrane (TM) region – which contains the channel pore – and a gigantic cytosolic “headpiece.” In this headpiece, cryo-EM studies have identified binding sites for native allosteric modulators of the channel function, such as calmodulin (CaM) (9) or FKBP12.0/12.6 (10). Leveraged synergies between cryo-EM, crystallography, and FRET have uncovered details of the structural mechanisms modulating the RyR function (1,11-13). Furthermore, recent breakthroughs in cryo-EM technologies have led to near-atomic high-resolution structures of the RyR1 and RyR2 (14,15). These include RyR co-structures with peptide toxins and small-molecule modulators, both physiologic and pharmacologic, that have been mapped to different regions of the channel complex (14,16-19).

To locate the *ent*-verticilide binding site within RyR2, here we used FLIM-detection of FRET within human RyR2 expressed in a stable HEK293 cell line. FLIM-FRET measurements from a set of donor-labeled FKBP12.6 (FKBP) variants (**Fig. 1**) to an acceptor-labeled *ent*-verticilide (**Fig. 2**) provided distance constraints for our trilateration method, which we used to locate the acceptor-labeled *ent*-verticilide probe within the cryo-EM map of RyR2.

**Fig 1.**
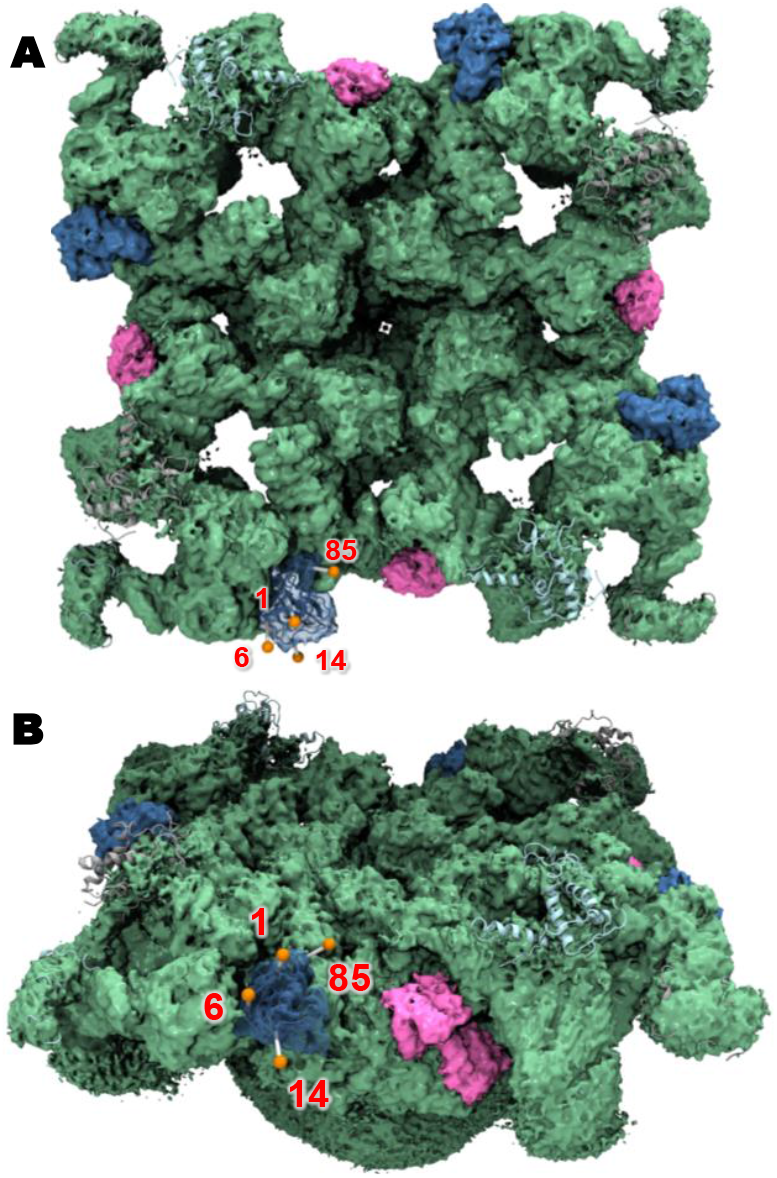
Molecular toolkit for RyR-specific FRET-based mapping of *ent*-verticilide. Cryo-EM structure of the FKBP12.6-RyR2 complex, with the effective positions of AF488 donor probes (orange spheres) attached at the indicated FKBP residues, as calculated via simulated annealing (1). A: Top view of the channel complex with FKBP and CaM represented in blue and pink, respectively. B: Tilted view of the RyR2 channel complex to provide a vantage point for the donor-location. In this study, we used FKBP labeled with donor at positions 1, 6, 14, or 85 (indicated by red numbers). Adapted from EMDB: 9833, PDB2: 6JI8.

**Fig 2.**
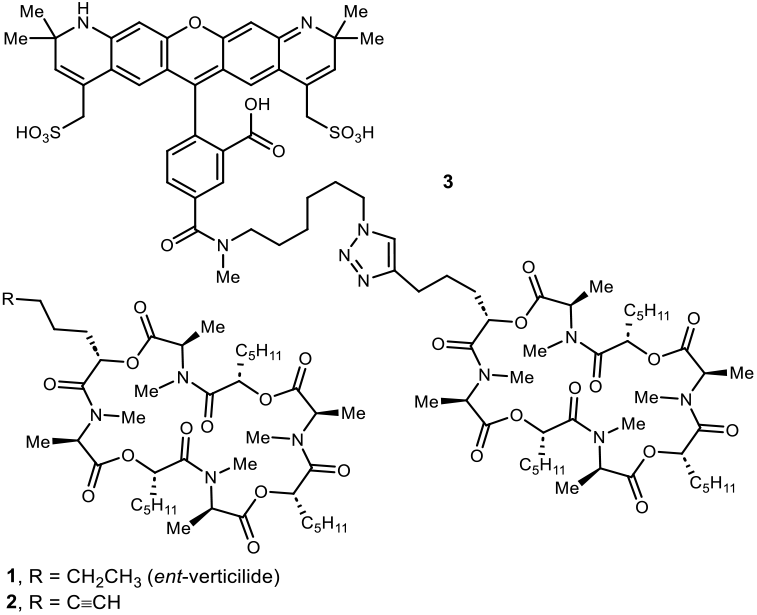
Chemical structures of *ent*-verticilide (**1**), alkyne analog (**2**), and AF-568-labeled analog of **2** (**3**).

## Results and Discussion

### Synthesis of AF568-*ent*-verticilide

FRET enables powerful approaches to resolve structural mechanisms of action within protein complexes. We have shown how distance analysis of FRET between donor-labeled sites of known locations and acceptor-labeled ligands can be used to trilaterate (map) RyR domains, accessory proteins or peptides binding regions within a protein complex such as the RyR channels (11,12). To use the same FRET-based method with *ent*-verticilide requires modifying it with an appropriate acceptor probe to obtain A-*ent*-verticilide. While the fluorescein analogue could be used, Alexa Fluor dyes exhibit reduced off-target binding (20). We chose Alexa Fluor 568 (AF568) because its optical properties are appropriate for pairing with our Alexa Fluor 488 (AF488)-labeled, RyR specific donor-targeting molecular constructs (1,21), a suite of AF488-labeled FKBPs (**Fig. 1B**).

AF568 is a versatile probe compatible with multiple types of fluorescence assays. Here, we paired AF568 as FRET acceptor for AF488 as FRET donor. In our experience, AF568 shows minimal non-specific binding effects compared with other red fluorophores, which is especially important for fluorescence labeling of relatively small molecules, such as *ent*-verticilide.

**Scheme 1.**
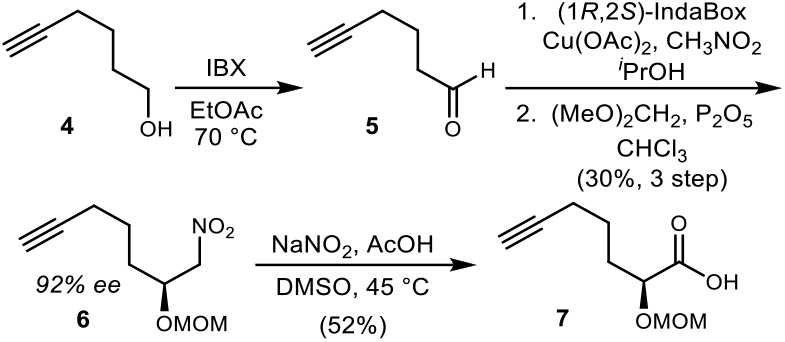
4-step synthesis of the unsaturated, *O*-protected α-hydroxy acid **7**.

The synthesis of AF568-labeled *ent*-verticilide (**Fig. 2** analog **3**) was planned from an alkyne-modified *ent*-verticilide (**Fig. 2**, analog **2**), providing a modest structural change that was not expected to affect activity against RyR2 (22). Two syntheses of *ent*-verticilide have been reported by us (7,23-26) but the oligomeric nature of **1** simplifies the didepsipeptide monomer synthesis and assembly. Non-oligomeric structures such as **2** (**Fig. 2**) require a de novo synthesis that preferably incorporates the modified residue at a late point in the synthesis. A terminal alkyne substituent for the α-oxy amide was selected for its potential to react in a final step with an azide-labeled fluorophore. This residue became a key synthetic challenge, requiring a suitably protected, enantioenriched α-hydroxy acid with a terminal alkyne chain (i.e. **7** in **Scheme 1**). While methods to access α-hydroxy acids are known (27-29), chemical enantioselective routes appropriate for this application are limited, owing to the chain length and need for a terminal alkyne. We elected to use an enantioselective Henry reaction (30-32).

Commercially available 5-hexynal (**4**) was oxidized to aldehyde **5** using IBX (**Scheme 1**) (20). The use of common alternative oxidants led to low yields and/or complex product mixtures. The aldehyde was then subjected to an enantioselective Henry reaction (29), followed by alcohol protection as the methoxymethylene (MOM) ether. This nitroalkane (**6**) was formed in 92% enantiomeric excess (ee) and 30% yield over 3 steps from the alcohol. Conversion of the terminal nitroalkane to a carboxylic acid was accomplished using Mioskowski’s oxidative variation of the Nef reaction (33), and the acid was isolated in 52% yield.

A trichloroethyl ester (TCE) was prepared with the expectation that this would provide a chemoselelective deprotection in a complex setting. This ester (**8**) was prepared in good yield by Steglich esterification (24,34), and then treated with boron trifluoride and an electrophile scavenger (pentamethyl benzene) (**Scheme 2**). The alcohol (**9**) was also used to prepare the alkyne-modified didepsipeptide **10** by Steglich esterification. *N*-Deprotection of **10** with HCl in dioxane provided the disubstituted amine needed for amidation with **11**. This was accomplished in two steps with 66% yield to furnish **12**. Selective deprotection of the ester proceeded smoothly with zinc dust, as did amine deprotection by HCl in dioxane, allowing cyclization using PyBop (35,36) to give the desired alkyne analog of *ent*-verticilide (**13**) (**Scheme 2**).

**Scheme 2.**
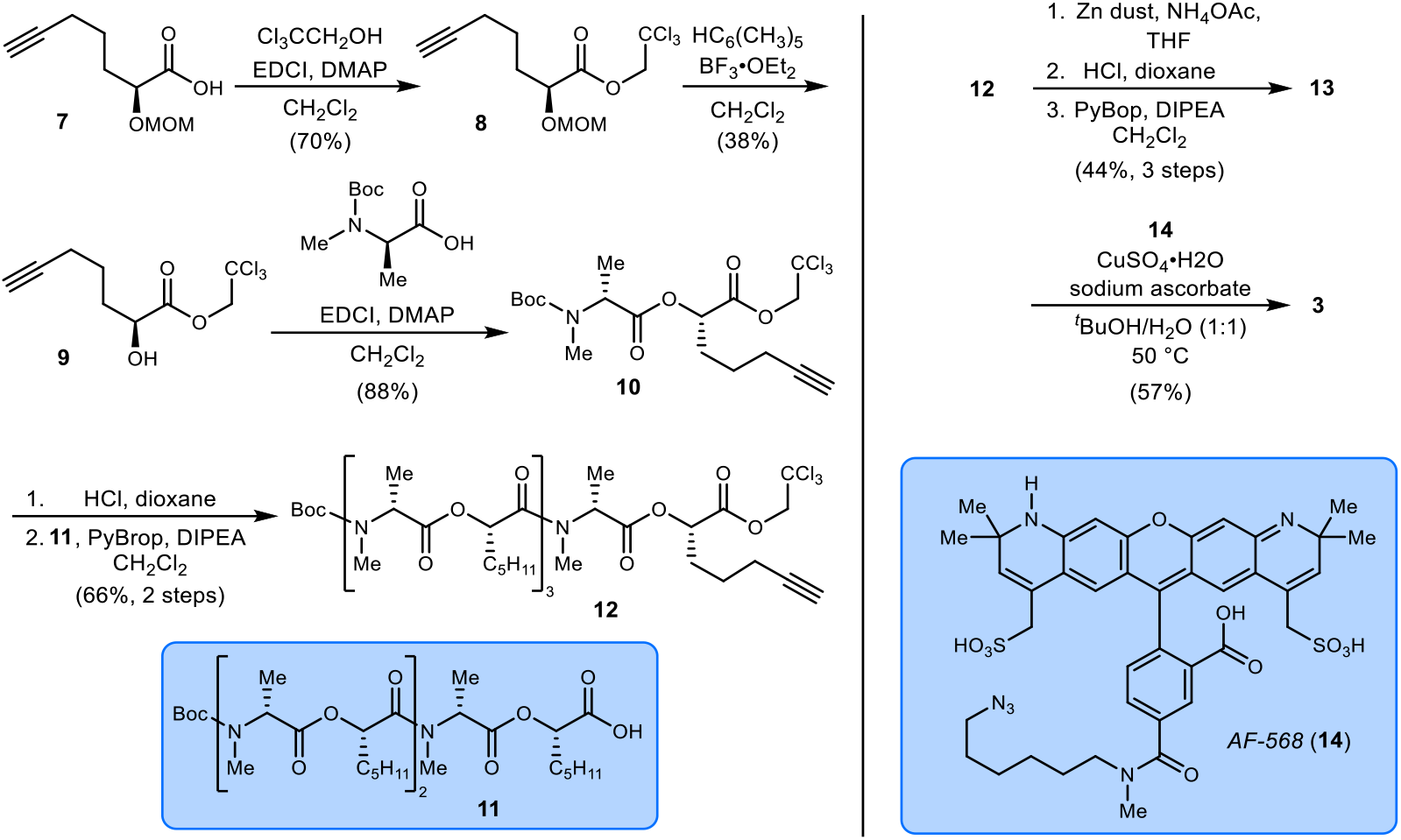
Synthesis of alkyne-labeled *ent*-verticilide (**2**) and its derivatization with AF568 to prepare **3**.

The success of this synthesis was due in part to the decision to form the ester bond in the didepsipeptide backbone instead of the amide (25). The amino acid can be incorporated in *N*-methylated form. The alternative would involve a central methylated amide bond and a terminal unprotected alcohol, a substrate expected to be highly prone to diketomorpholine formation (7,37).

The alkyne analog (**13**) was evaluated in the Ca^2+^ sparks assay, revealing inhibition similar to *ent*-verticilide (data not shown). This finding established that one of the four alkyl side chains in the cyclic oligomeric parent *ent*-verticilide can be modified, at least subtly, without a significant loss of activity. Alkyne **13** was then subjected to a Huisgen cycloaddition using commercially available AF568 azide (**Scheme 2**) (38-40). While the reaction proceeded smoothly to provide the desired product (reaction was monitored by LC/MS for both appearance of product mass, and consumption of starting materials.) the workup and purification of this fluorescent analogue proved to be difficult owing to the solubility of **3** in water. Fortunately, it could be salted-out (sodium chloride). The crude material was then purified via reverse phase preparatory HPLC. Following lyophilization, analysis by ^1^H NMR provided support of the assignment as in **3**. When tested in the Ca^2+^ spark assay, AF568-*ent*-verticilide (**3**) was similarly potent to *ent*-verticilide (**Fig. 3A**).

**Fig 3.**
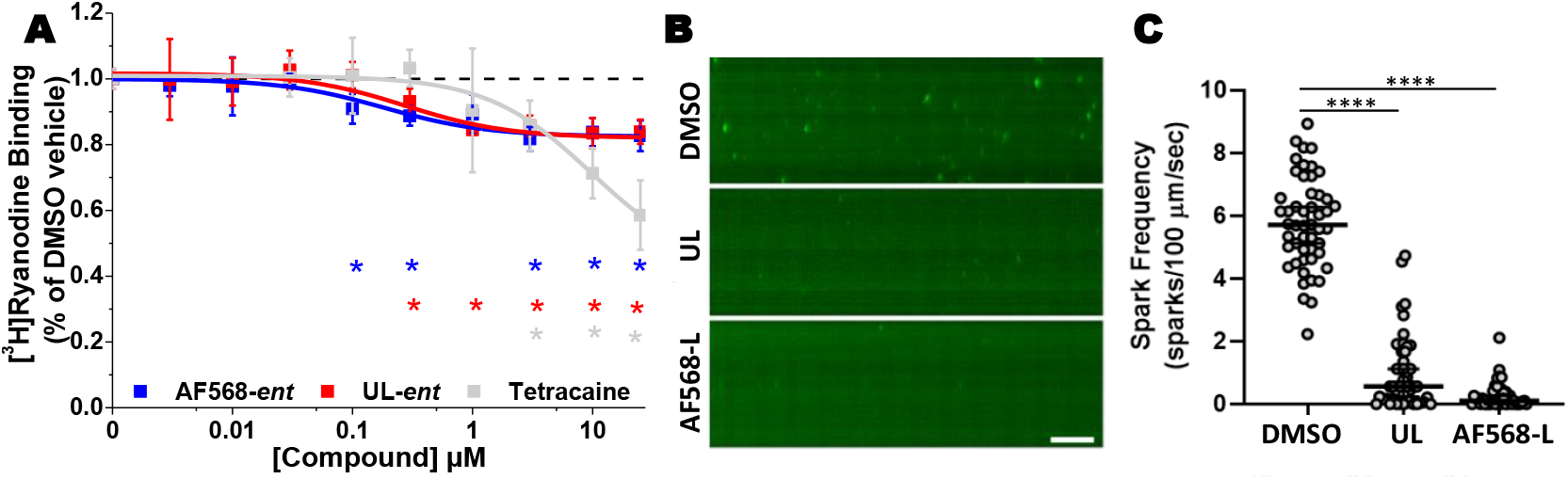
Functional validation of AF568-labeled *ent*-vert via RyR2-specific functional assays. (A) [^3^H]ryanodine binding measurements of AF568-labeled (AF568-L) and unlabeled (UL) *ent*-vert dose response of RyR2 function. Measurements were carried out using cardiac SR vesicles (RyR2) from porcine tissue in the presence of 100 nM [Ca^+2^], as described in Methods. Error bars indicate standard error (N=6). Statistical significance was determined via student’s t-test (2-tailed, paired); **\***p<0.05. (B) Imaging of Ca^2+^ sparks in mouse ventricular myocytes: representative line scans from permeabilized cardiomyocytes showing illustrating the effect of *ent*-vert unlabeled (UL) and acceptor-labeled (AF568-L) on Ca^2+^ sparks, relative to a vehicle control (DMS). Scale bar = 0.5 s; (C) Ca^2+^ spark frequency N(DMSO) = 50 cells from 3 mice; N(UL) = 48 cells from 3 mice; N(AF568-L) = 39 cells from 3 mice. Statistical significance was determined via hierarchical clustering; ****p<0.0001 vs DMSO.

### Validation of AF568-*ent*-verticilide using functional measures of RyR2

To determine whether the *ent*-verticilide action against RyR2 is preserved after chemical modification AF568 we used biochemical and myocyte assessments of RyR2 function.

First, we measured [^3^H]ryanodine binding to isolated cardiac SR (containing RyR2), which is a widely-used assay to quantitate channel opening in a population of RyRs. We assessed the concentration-dependent effects of the AF568-derivatized *ent*-verticilide analog vs. unmodified *ent*-verticilide, on the [^3^H]ryanodine binding to RyR2 (**Fig. 3A**). We carried out these assays with labeled and unlabeled [*ent*-verticilide] ranging from 30 nM to 30 μM (**Fig. 3A**), to determine their effect on [^3^H]ryanodine binding to isolated cardiac SR membranes, in 100 nM (diastolic) Ca^2+^. For unlabeled *ent*-verticilide, we observed an RyR2 inhibition profile similar to our previous report (7), with submicromolar EC_50_ and plateauing at about 20% inhibition. The AF568-*ent*-verticilide analogue trends to a slightly lower EC_50_ (higher potency) but a similar level of inhibition as the unlabeled compound. As a control, we measured response to tetracaine, which shows the expected inhibition profile. Thus, the [^3^H]ryanodine assays suggest that the AF568 labeling of *ent*-verticilide does not alter its properties against channel opening in the resting state of RyR2.

We next examined Ca^2+^-spark frequency in isolated murine cardiomyocytes treated with AF568-derivatized *ent*-verticilide analog vs unmodified *ent*-verticilide (**Fig. 3B,C**). Ca^2+^ spark frequency is routinely used as a measure of RyR2-specific Ca^2+^ leak in the cardiomyocyte. Pathological modifications of the RyR2 channel complex, such as hyperphosphorylation, oxidation or arrhythmogenic mutations, are associated with increased spark frequency (41-43). Conversely, interventions that reduce spark frequency are regarded as anti-arrhythmic because they reduce the inappropriate leak of Ca^2+^ during diastole (41,42,44). RyR2-mediated Ca^2+^ sparks were recorded from permeabilized murine Casq2^-/-^ cardiomyocytes to determine the effect of the attachment of AF568 to *ent*-verticilide. Ca^2+^ spark frequency was reduced to a similar extent by *ent*-verticilide and AF568-*ent*-verticlide. Taken together, these results indicate that attachment of AF568 to *ent*-verticilide did not disrupt its binding to, or inhibition of, RyR2.

### FLIM-FRET between AF488-FKBPs and AF568-*ent*-verticilide in HEK293 cells expressing human RyR2

AF568-*ent*-verticilide can be evaluated for binding and location within the RyR2 structure using our previously reported FRET-based trilateration methods, which were demonstrated on myocyte-expressed RyR2 (1) and HEK293-expressed RyR1 (11,12,45).

To describe *ent*-verticilide binding within RyR2, we took advantage of our previously reported stable expression of the *human* RyR2 isoform in an HEK293 cell line (hRyR2-HEK293). For this analysis, we used four donor-labeled FKBPs to carry out FLIM measurements of FRET to AF568-labeled *ent*-verticilide in hRyR2-HEK293 cells (**Fig. 4**). With these measurements, we aimed to resolve distance relationships between donor probes at known locations on RyR2-associated FKBP and the acceptor probe on *ent*-verticilide, to map *ent*-verticilide binding within the cryo-EM structure of RyR2.

**Fig 4.**
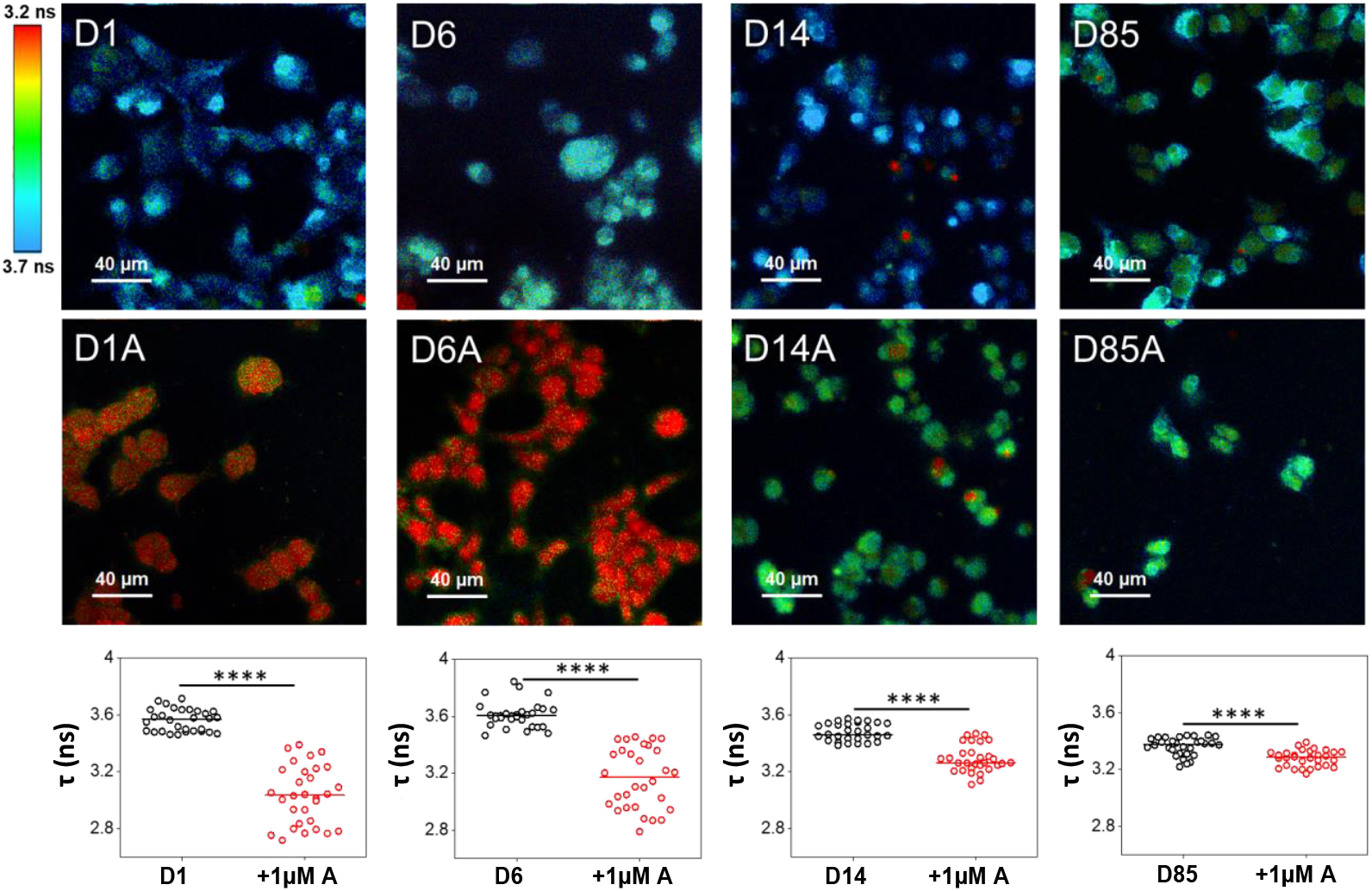
FLIM-FRET Imaging and Spectroscopy. The top panels show representative images of the donor-only imaging, for donor at FKBP position D1, D6, D14, or D85, from left to right. The middle panels show the corresponding samples after addition of 1 μM AF568-*ent*-verticilide. The bottom panels summarize the τ values for the donor-only, donor plus 0.2 μM AF568-*ent*-verticilide, and 1 μM AF568-*ent*-verticilide as indicated. All donors showed shorter τ in the presence of the acceptor as is indicated by the shift toward warmer colors in the FLIM images. The same shift toward shorter amplitude-weighted average τ was detected for single-cells TCSPC measurements with focused laser beam (bottom panels). The significance of the decrease in τ was determined using two-sample T-test with significance level of 0.05. *p<0.05; **p<0.01: ***p<0.001; ****p<0.0001.

For this experiment, we used permeabilized hRyR2-HEK293 cells pre-incubated with AF488-labeled FKBP at single-Cys residues substituted at positions 1, 6, 14, and 85. For each AF488-FKBP, we first acquired donor (D)-only images (**Fig. 4**, top panels), observing FLTs (τ) for donor attached at FKBP residues 1, 6, 14, or 85 (Table 1, τ_D_). Addition of a saturating concentration of AF568-*ent*-verticilide acceptor (A; 1 μM final concentration; **Fig. 4**, middle panels) led to shorter donor τ (Table 1, τ_D+A_). The shortened τ values suggest that the acceptor binds within FRET range (i.e., <100Å) from each donor probe. Combined, the donor-only (D) and donor-acceptor (DA) data yield FRET efficiencies (*E*) ranging from 0.02 to 0.15 (Table 1). A cursory evaluation of the FRET values suggests that the acceptor is closer to the donors attached at FKBP positions 1 and 6 (higher *E*) than to the donors attached at 14 and 85 (lower *E*).

**Table 1.** FLT-detected FRET measurements and distance relationships between donor-labeled FKBP and acceptor-labeled *ent*-verticilide bound to hRyR2-HEK293 cells.*

| D site<sup>a</sup> | $\langle\tau_D\rangle^b$ | $\langle\tau_{D+A}\rangle^c$ | $E^d$ | $R$ (Å)<sup>e</sup> |
| --- | --- | --- | --- | --- |
| 1 | 3.56(.01)<sup>f</sup> | 3.03(.04) | 0.15(.002) | 83(2) |
| 6 | 3.61(.02) | 3.17(.04) | 0.12(.002) | 86(2) |
| 14 | 3.48(.01) | 3.29(.02) | 0.05(.001) | 100(1) |
| 85 | 3.36(.01) | 3.28(.01) | 0.02(.001) | 114(1) |

\*Analysis of **Fig. 4** FLT data.
<sup>a</sup>Location of single-Cys substitution in the sequence of null-Cys FKBP. Each single-Cys FKBP variant was labeled with AF488;
<sup>b</sup>Amplitude-weighted average FLT values in donor-only samples;
<sup>c</sup>Amplitude-weighted average FLT values in donor-plus-acceptor samples;
<sup>d</sup>FRET efficiency, $E$ , was calculated according to Eq. (1);
<sup>e</sup>Distances $R$ were calculated according to Eq. (2);
<sup>f</sup>Standard error is indicated between parentheses;

This striking pattern of higher FRET for donors that are proximal to the corners of the RyR cytosolic portion and lower FRET for donors that are proximal to the center of each face of the RyR cytosolic portion suggests an eccentric binding location for A-*ent*-verticilide, away from the channel pore that is centrally located, deep in the transmembrane domain of the RyRs. This supports the notion that *ent*-verticilide is an allosteric inhibitor of RyR2 rather than an orthosteric channel blocker. This is consistent with the fractional inhibition exerted by *ent*-verticilide in [^3^H]ryanodine binding assays using cardiac SR samples (**Fig. 3A**).

### Mapping the AF568-*ent*-verticilide location within the RyR2 structure

For a more precise determination of the A-*ent*-verticilide location, we used these four distances with our previously reported trilateration protocol (1). Trilateration is a method that uses distances from multiple known positions to determine the position of a point of interest. As an example, a broadly used application of trilateration is GPS positioning. A minimum of four distances from a FRET donor probe is necessary to trilaterate the location of an acceptor probe in 3D space. With one distance measurement the acceptor would be located on the surface of a sphere. Adding one more distance measurement from another donor would locate the acceptor at the intersection of two spheres, which is a toroid shape. By adding a third distance measurement we get the intersection of three spheres which has two solutions (i.e., two loci). By adding a fourth measurement one could pinpoint a specific locus in space. This represents an ideal situation where distances can be determined with high precision from well-spaced, precisely determined donor positions. In experimental reality, results tend to be more ambiguous, and even with extra distances multiple loci might result. Unique locations can still be inferred by integration of other experimental constraints (1).

From the time-correlated single photon counting (TCSPC)-FRET measurements, we determined the distances, *R*, separating each donor probe attached to FKBP from the acceptor probe attached to *ent*-verticilide, calculated according to Eq. (2) (Table 1). Trilateration calculations were based on methods we previously described (1). For this process, the effective donor locations were determined from simulated annealing calculations of AF488 bound to FKBP in an RyR2 cryo-EM high-resolution co-structure with bound FKBP and CaM (46). Distances (*R*) from donor to acceptor labels were calculated based on the average FRET efficiencies (*E*) determined from TCSPC-FRET experiments and the Förster distance characteristic for AF488-AF568 donor-acceptor pair (*R*_0_ = 62 Å).

Trilateration based on FRET-derived distances from donor-labeled FKBP identifies two possible binding locations (loci L1 and L2 in **Fig. 5**) for A-*ent*-verticilide on RyR2. These loci represent the location of the acceptor fluorophore, which is predicted to extend out into the solvent about 10-12 Å out from the *ent*-verticilide binding site. The first locus (L1 in **Fig. 5**) is found at the periphery of the channel complex cytoplasmic portion, within the P1 domain (aka Ry12 domain). This location is far (>100 Å) from the channel pore whose opening behavior *ent*-verticilide modulates. The second locus (L2 in **Fig. 5**) is found at the interface formed between the SPRY3 domain of one RyR2 protomer and the upper part of the HD1 domain of the next RyR2 protomer of the channel assembly. This location is on the other side of the HD1 domain from the CaM binding location on RyR2. In the L2 location, *ent*-verticilide may affect protomer-protomer interactions within the RyR2 channel complex. Although both L1 and L2 are quite far from the channel pore, modulation of RyR opening is known to involve long-range structural pathways of long-distance allostery (47,48).

**Fig 5.**
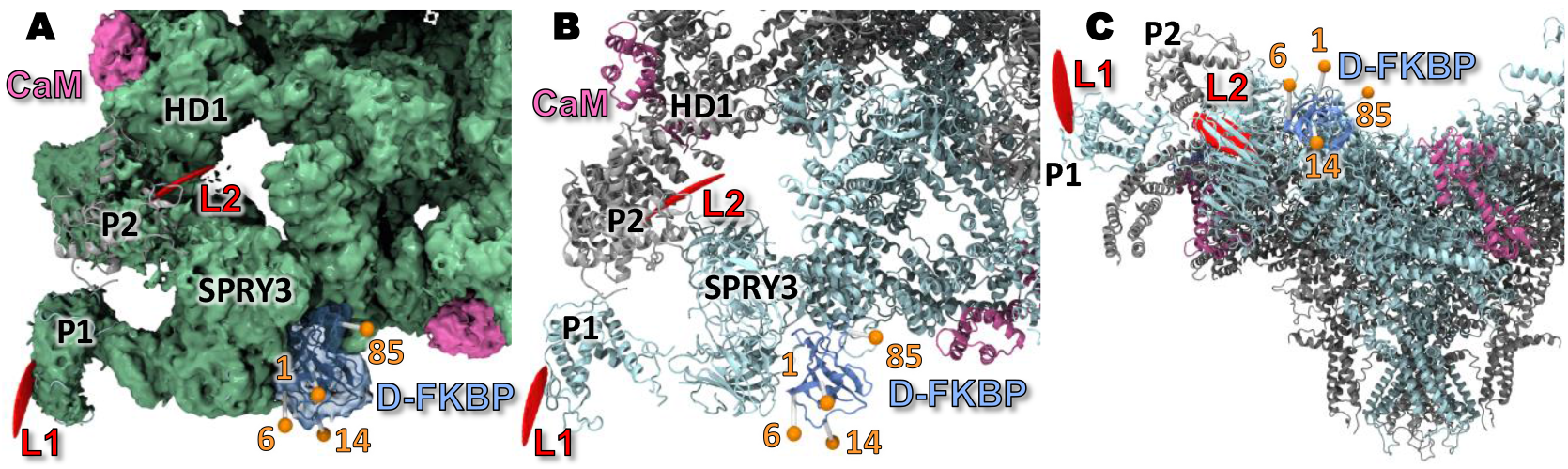
Mapping of the A-ent-verticilide binding within the cryo-EM structure of RyR2. Trilateration based on FRET-derived distances from donor-labeled FKBP identifies two possible binding locations (L1 and L2) of A-*ent*-verticilide within RyR2, indicated in red in the cryo-EM density map (panel A) and corresponding ribbon structures (panels B and C). Effective donor locations (orange spheres) were determined *in silico* via simulated annealing of AF488 bound to FKBP in an RyR2 structure with bound FKBP and CaM (6JI8) (46). Donor-acceptor distances were calculated from FRET efficiencies (Table 1). These distances were used to calculate locations of the acceptor fluorophore, which is predicted to be 10-12 Å from the *ent*-verticilide binding site. L1 is adjacent to the P1 domain as can be seen in the top-view density map (panel A) and ribbon structure (panel B). L2 is found near the inter-subunit interface formed between the SPRY3 domain of one RyR2 protomer and the HD1 domain of the adjacent protomer in the channel complex (panels A and B). In panel C, binding location L2 can be observed behind the SPRY3 domain.

### Relationship to other small-molecule binders

Multiple small-molecule modulators of RyRs have been investigated through cryo-EM. Two of these directly map to the transmembrane region and can be considered channel activators: diamide insecticides bind to the pseudo-voltage sensing domain and trigger pore opening (16), whereas ryanodine binds directly within the ion conduction pathway in the pore-forming domain (14). Although ryanodine can block RyRs at high concentrations, at lower concentrations it keeps the channel in a long-lasting subconducting state. Together with ryanodine’s slow binding kinetics, this precludes its use as a therapeutic agent. Close to the transmembrane region are binding sites for Ca^2+^, ATP, and caffeine, which also activate the channel. The binding site for ATP can also be occupied by other adenine derivatives, including cAMP (18).

However, one proposed inhibitory class of molecules, known as rycals, were recently observed bound to the P1 domain, in complex with an ATP molecule (17,49). Interestingly, one of our proposed sites for the *ent*-verticilide (L1 in **Fig. 5**) also resides within the P1 domain, indicating the potential importance of this domain in regulating pore opening. Of note, the P1 domain is implicated in inter-RyR interactions (50), and modulating the conformation of this domain may thus affect such interactions. Whether rycals and *ent*-verticillide act through the same mechanism remains to be tested.

A second possible site for *ent*-verticillide (L2 in **Fig. 5**) is at the interface between the SPRY domains and the HD1 domain. The HD1 domain also forms the binding site for CaM, and affecting its conformation, either through binding of CaM or through the introduction of disease-causing mutations, has been linked to channel gating (46,51). The SPRY-HD1 interface was also found to be labile in the presence of calcins, a class of scorpion peptides that keep the pore in an open conformation (19). The SPRY domains themselves are also the target for disease-associated mutations (52). Thus, molecules binding to this region have the potential to affect channel gating.

## Methods

### Isolation of sarcoplasmic reticulum vesicles

Crude sarcoplasmic reticulum membrane vesicles were isolated from porcine cardiac left ventricle tissue by differential centrifugation of homogenized tissue, as established previously (53). These samples were flash-frozen and stored at -80ºC until needed for experiments.

### Stripping of endogenous CaM

Isolated cardiac SR vesicles were incubated with, 300 nM CaM binding peptide, 0.1 μM CaCl2, 20 mM PIPES, 150 mM KCl, 5mM GSH, 0.1 mg/mL BSA, 1 μg/mL aprotininin, 1μg/mL leupeptin, and 1 μM DTT for 30 min at 37°C. Samples were then spun at 110,000x*g* for 25 min at 4°C and resuspended to at final concentration of 15 mg/mL in 20 mM PIPES, 150 mM KCl, 5mM GSH, 0.1 mg/mL BSA, 1 μg/mL aprotinin, 1μg/mL leupeptin, and 1 μM DTT. These CaM-stripped vesicles were immediately used for [^3^H]ryanodine binding protocols.

### [^3^H]ryanodine binding to SR vesicles and data analysis

In 96-well plates, cardiac SR membranes (CSR, 3 mg/mL) were pre-incubated with 1% DMSO or AF568-labeled or unlabeled *ent*-verticilide, for 30 min, at 22°C, in a solution containing 150 mM KCl, 5 mM GSH, 1 μg/mL Aprotinin/Leupeptin, 1 mM EGTA, and 238 μM CaCl_2_ (as determined by MaxChelator to yield 100 nM free Ca^2+^), 0.1 mg/mL of BSA, 5mM Na_2_ATP and 20 mM K-PIPES (pH 7.0). Non-specific [^3^H]ryanodine binding to SR was assessed by addition of 40 μM non-radioactively labeled (“cold”) ryanodine. Maximal [^3^H]ryanodine binding was assessed by addition of 5 mM adenylyl-imidodiphosphate (AMP-PNP), supplemented with 20 mM caffeine. These control samples were each loaded over four wells per plate. Binding of [^3^H]ryanodine (7.5 nM) was determined following a 3 h incubation at 37°C and filtration through grade GF/B glass microfiber filters (Brandel Inc., Gaithersburg, MD, US) using a 96-channel harvester (Brandel M-96). Filters were immersed in 4 mL of Ecolite scintillation cocktail for 24 h before [^3^H] counting in a Perkin-Elmer Tri-Carb 4810.

### Ca^2+^ imaging in cardiomyocytes

The care and use of animals followed established guidelines and was approved by the Animal Care and Use Committee of Vanderbilt University Medical Center (Protocol number M1900057-00). Mouse cardiomyocytes were isolated and prepared as previously described (7). Cells were adhered to laminin-coated #1 coverglass and then permeabilized using saponin with an internal solution designed to promote RyR2-mediated Ca^2+^ sparks (7). The internal solution contained 14 μg/mL Fluo-4 to detect Ca^2+^ and 25 μM DMSO (vehicle), *ent*-verticilide, or AF568-*ent*-verticilide. Ca^2+^ sparks were recorded after a 10-minute incubation with the test compound, and line scan recordings were analyzed using the SparkMaster plugin for ImageJ.

### Confocal imaging of hRyR2-HEK293 cells

A stable, inducible Flp-In T-Rex-293 cell line expressing human RyR2 was generated using the Flp-In T-REx Core Kit (Invitrogen, USA) as described before (54). Specifically, Flp-In T-REx-293 cells were co-transfected with the pOG44 vector encoding the Flp recombinase and the expression vector pcDNA5/FRT/TO containing the unlabeled human RyR2 cDNA. The growth medium was replaced 48 h post-transfection with a selection medium containing 100 μg/ml hygromycin. The hygromycin-resistant cell foci were selected and expanded. Expression of human RyR2 in the stable cell line was verified by western blot analysis 48 hours post-induction of recombinant RyR2 expression with 1 μg/ml tetracycline. Flp-In T-Rex-293 human RyR2 stable line cells were plated on 35 mm plastic dishes and grown in high-glucose DMEM supplemented with 100 U/ml penicillin, 100 mg/ml streptomycin, and 10% fetal bovine serum at 5% CO_2_ and 37°C. After 24 hours, 1 μg/ml of tetracycline was added to induce the expression of RyR2 for another 48 hours. Before the experiment, the cells were transferred to laminin-coated Lab-Tex chamber slides (ThermoFisher Scientific, USA) for the corresponding imaging experiments.

Immediately before the experiment, the cells were washed once in phosphate-buffered saline (PBS) and subsequently washed with low-Ca^2+^ relaxation solution (120 mM potassium aspartate, 15 mM KCl, 5 mM KH_2_PO_4_, 0.75 mM MgCl_2_, 2% dextran, 5 mM ATP, 20 mM HEPES, and 2 mM EGTA, pH 7.2). Cells were permeabilized with 0.005% saponin solution for 5 minutes and washed twice with high-Ca^2+^ solution composed of 120 mM potassium aspartate, 15 mM KCl, 5 mM KH_2_PO_4_, 0.75 mM MgCl_2_, 2% dextran, 5 mM, 20 mM HEPES, 2 mM EGTA, and 1.7 mM CaCl_2_, pH 7.2. The free [Ca^2+^]_i_ concentration was estimated as 2 μM (55).

TCSPC experiments were performed as previously described (56-58). TCSPC histograms for donors without the acceptor were acquired for permeabilized RyR2-expressing cells that were incubated for 15 min at room temperature in the presence of FKBP12.6 labeled with Alexa Fluor 488 at positions 1, 6, 14, or 85. Subsequently, 1 μM Alexa Fluor 568-labeled *ent*-verticilide was added, and incubated for 15 min and TCSPC histograms were acquired for the donor in the presence of the acceptor. Donor fluorescence in the cell was excited by a supercontinuum laser beam (Fianium). Donor (AF568-FKBP) signal was acquired using a 482/18 nm bandpass filter in the excitation channel. Fluorescence emission was detected using a 525/50 nm bandpass filter in the emission channel. After cell selection using the defocused laser beam, the defocusing lens was removed from the light path, the laser intensity was attenuated with a neutral density filter (OD = 0.5), and the focused laser was placed on the cell membrane, yielding a count rate of 100,000 photons/s rate. Under those conditions, we observed less than 10% photobleaching during the 60-s acquisition. Fluorescence was detected through a 1.2 numerical aperture water-immersion objective with a PMA hybrid detector (PicoQuant, Germany) connected to a single photon-counting module (HydraHarp 300, PicoQuant, Germany) with a time-channel width of 16 ps. Fluorescence decay histograms from 10 cells from a single independent transfection were analyzed in SymPho Time 64 software with TCSPC fitting tool. All donors showed two-exponential decay with relative amplitude of the second component accounting for 11-16%. Therefore, the donor alone FLTs (τ_D_) were determined from 2-exponential fits. The fluorescence decays of donor in the presence of saturating acceptor required a 3-exponential fit. Therefore, the donor-plus-acceptor FLTs (τ_DA_) were determined from 3-exponential fits. For both donor-only and donor-plus-acceptor decays, the amplitude-weighted average FLTs (<τ_D_> and <τ_DA_>, respectively) were used to calculate the FRET efficiency (*E*) according to

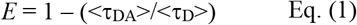

From *E* and the Förster distance, *R*_0_ = 62Å, characteristic to the AF488-AF568 donor-acceptor pair, we calculated the distances separating each of the FKBP12.6-attached donor from the *ent*-verticilide-attached AF568, according to:

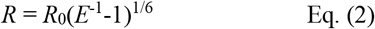

In the calculations of *R*, we assumed the orientation factor κ^2^ = 2/3, as previously discussed (1). Values for τ_D_, τ_DA_, *E*, and *R* are reported in Table 1.

### Trilateration procedure

With FRET-derived distances as input, we used a previously described trilateration method to determine AF568-*ent*-verticilide loci as candidate binding sites (1). We used the 3.6-Å RyR2 structure in the closed state (PDB ID: 6IJ8) and the corresponding cryo-EM density map (EMDB ID: 9833) to calculate the donor probe locations. This included two new donor-FKBPs (D1 and D6) that were added since the original publication. A simulated annealing protocol was used as described earlier to determine an ensemble of probe locations. Using the assumption that the fluorescence probe shows fast dynamics in relation to the florescence life-time (59), the average center position for the florescent probe was calculated from the simulated annealing.

From the simulated-annealing calculations above, we obtained coordinates of the average probe locations, which we use as effective donor locations for trilateration. We used effective AF488 locations calculated for the four FKBP attachment sites (1, 6, 14, and 85), and distances, *R*, calculated from FRET efficiency values, *E*, to determine a location in space corresponding to the acceptor attached to *ent*-verticilide. The other input value required in the trilateration method is a range around the calculated *R*. Based on prior results using other systems, we estimated this range to be 10% of *R*. Using a proportional estimation is a reasonable alternative because uncertainties increase as *R* lengthens beyond *R*_0_ (1).

The normalized sum of the distance ranges from each donor site is generated as a map in 3D space that describes the probability of the acceptor location. This volumetric map is saved in MRC/CCP4-format, which can be displayed using common molecular graphics software. Figures were created using VMD (60).

## Statistical analysis

Errors are reported as the standard error of the mean (SE), except when noted. Statistical significance was determined by Student’s t-test, hierarchical clustering or one-way ANOVA followed by Tukey’s post-hoc test, as indicated, where p<0.05 was considered significant. EC_50_ values were derived from the fits to Hill equations.

## Safety Statement

No unexpected or unusually high safety hazards were encountered.

## Data availability

All data are contained within the manuscript.

## Supporting information

[Link]

## Acknowledgements

This work was supported by American Heart Association Postdoctoral Fellowship 830562 (to J. S.) and the National Institute of Health grants F31 HL151125 (to A.N.S.), R01 HL092097 and HL138539 (to R.L.C), HL151223 (to J.N.J., B.C.K., and R.L.C.), HL158649 (to S.L.R., and R.L.C.), and HL151990 (to A.V.Z.). The content is solely the responsibility of the authors and does not necessarily represent the official views of the National Institutes of Health.

