## Supplemental Information for "RyR2 Binding of an Antiarrhythmic Cyclic Depsipeptide Mapped Using Confocal Fluorescence Lifetime Detection of FRET"

**Contents** **SI-X**

### Experimental Section

**General Information:** Glassware was flame-dried under vacuum for all non-aqueous reactions. All reagents and solvents were commercial grade and purified prior to use when necessary. Benzene and dichloromethane ($CH_2Cl_2$) were dried by passage through a column of activated alumina as described by Grubbs.[1] Flash column chromatography was performed using Sorbent Technologies 230-400 mesh silica gel with solvent systems indicated. Analytical thin layer column chromatography was performed using Sorbent Technologies 250 µm glass backed UV254 silica gel plates and were visualized by fluorescence upon 250 nm radiation and/or the by use of ceric ammonium molybdate (CAM), phosphomolybdic acid (PMA), or potassium permanganate (KMnO4). Solvent removal was affected by rotary evaporation under vacuum (~ 25-40 mm Hg). All extracts were dried with $MgSO_4$ or $Na_2SO_4$ unless otherwise noted. Preparative HPLC was performed on an Agilent 1260 system (column: Zorbax Eclipse XDB-C18; 21.2 mm x 150 mm, 5 µm, flow rate 8 mL/min) with 210 nm monitoring wavelength and acetonitrile/water (+0.1% TFA) gradient as indicated. Nuclear magnetic resonance spectra (NMR) were acquired on a Bruker AV-400 (400 MHz), Bruker DRX500 (500 MHz), or Bruker AV II-600 (600 MHz) instrument. Chemical shifts are measured relative to residual solvent peaks as an internal standard set to δ 7.26 and δ 77.16 ($CDCl_3$), unless otherwise specified. Mass spectra were recorded by use of chemical ionization (CI), electron impact ionization (EI), or electro-spray ionization (ESI) on a high resolution Thermo Electron Corporation MAT 95XP-Trap by the Indiana University Mass Spectrometry Facility or on a TQ-Orbitrap 3 XL Penn or Orbitrap 2 Classic FPG in the Vanderbilt Mass Spectrometry Core Laboratory. IR spectra were recorded on a Nicolet Avatar 360 spectrophotometer and are reported in wavenumbers ($cm^{-1}$) as neat films on a NaCl plate (transmission). Melting points were measured using an OptiMelt automated melting point system (Stanford Research Systems) and are not corrected. Chiral HPLC analysis was conducted on an Agilent 1100 series Infinity instrument using the designated ChiralPak column. Optical rotations were measured on either a Jasco P-2000 polarimeter or an AUTOPOL III (Rudolph Research Analytical) polarimeter.

**Purity Statement:** No in vivo work is reported. Although each final product was purified by preparative HPLC for in vitro assay, and detected by UV (210 nm), the compounds reported lack a good chromophore. However, the use of HPLC in combination with NMR analysis led us to judge compounds to be a minimum of 90% pure, likely >95%. Images of NMR spectra of all new compounds are provided.

**Safety:** No unexpected or unusually high safety hazards were encountered

---

[1] Pangborn, A. B.; Giardello, M. A.; Grubbs, R. H.; Rosen, R. K.; Timmers, F. J. *Organometallics* **1996**, *15*, 1518.

### Experimental and Characterization Data for Reported Compounds

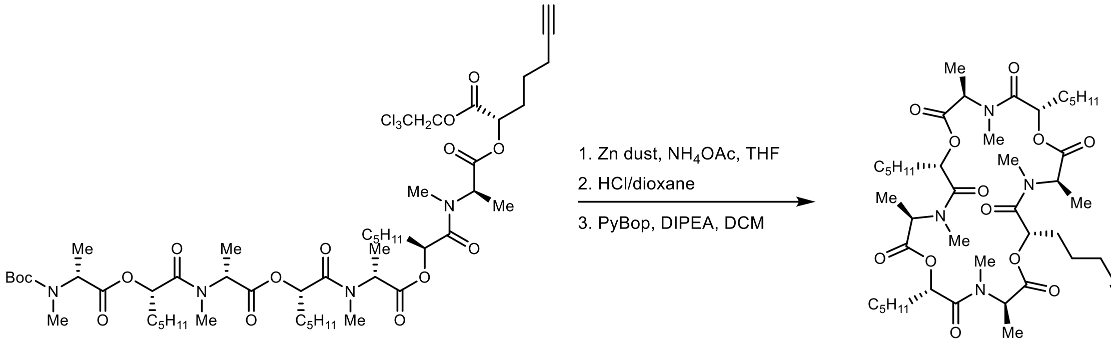

**(3*R*,6*S*,9*R*,12*S*,15*R*,18*S*,21*R*,24*S*)-3,4,9,10,15,16,21,22-Octamethyl-6-(pent-4-yn-1-yl)-12,18,24-tripentyl-1,7,13,19-tetraoxa-4,10,16,22-tetraazacyclotetracosan-2,5,8,11,14,17,20,23-octaone (*ent*-2).** A round-bottom flask was charged with the depsipeptide (26.0 mg, 23.7 µmol), dissolved in THF (1.47 mL) and 1 M $NH_4OAc$ (260 µL), and treated with zinc dust (77 mg). The reaction was allowed to stir for 2 h and then the crude reaction mixture was filtered through Celite. To the crude material was added 4 M HCl in dioxane (400 µL) and the reaction mixture was allowed to stir for 30 min. The reaction mixture was concentrated and added to a flame-dried round-bottom flask. DCM (5.1 mL) was added and the reaction was cooled to 0 °C. Once at 0 °C, DIPEA (9.6 µL, 56 µmol) and PyBrop (13.9 mg, 26.8 µmol) were added. The reaction was stirred at 0 °C for 1 h, then allowed to warm to ambient temperature and stir for an additional 1 h. The reaction mixture was poured into cold 10% aq citric acid and extracted with DCM. The combined organic layers were washed with satd aq $NaHCO_3$ and brine, and then dried and concentrated. Preparative HPLC (5 – 95% aqueous acetonitrile, 210 nm, flow rate: 8 mL/min, $R_t$ = 21 m) afforded the 24-membered macrocycle (8.9 mg, 44%) as a colorless oil. $[\alpha]_D^{24}$ +34 (*c* 0.51, $CHCl_3$); $R_f$ = 0.18 (4% MeOH/DCM); IR (film) 3284, 2931, 2862, 1742, 1661, 1461, 1414, 1317, 1199, 1085 $cm^{-1}$; $^1H$ NMR (600 MHz, $CDCl_3$) This compound exists in multiple conformations, causing significant peak overlap. Refer to the image of the $^1H$ NMR spectrum; $^{13}C$ NMR (150 MHz, $CDCl_3$) This compound exists in multiple conformations, causing significant peak overlap. Refer to the image of the $^{13}C$ NMR spectrum; HRMS (EI): Exact mass calcd for $C_{44}H_{72}N_4NaO_{12}$ $[M+Na]^+$ 871.5039, found 871.5051. ANS-3-270.

**(3*S*,6*R*,9*S*,12*R*,15*S*,18*R*,21*S*,24*R*)-3,4,9,10,15,16,21,22-Octamethyl-6-(pent-4-yn-1-yl)-12,18,24-tripentyl-1,7,13,19-tetraoxa-4,10,16,22-tetraazacyclotetracosan-2,5,8,11,14,17,20,23-octaone (2).** Prepared following an identical procedure as *ent*-**2**. Preparative HPLC (5 – 95% aqueous acetonitrile, 210 nm, flow rate: 8 mL/min, $R_t$ = 21 m) afforded the 24-membered macrocycle with spectroscopic data identical to its enantiomer, except $[\alpha]_D^{24}$ -33 (*c* 0.52, $CHCl_3$). ANS-4-104.

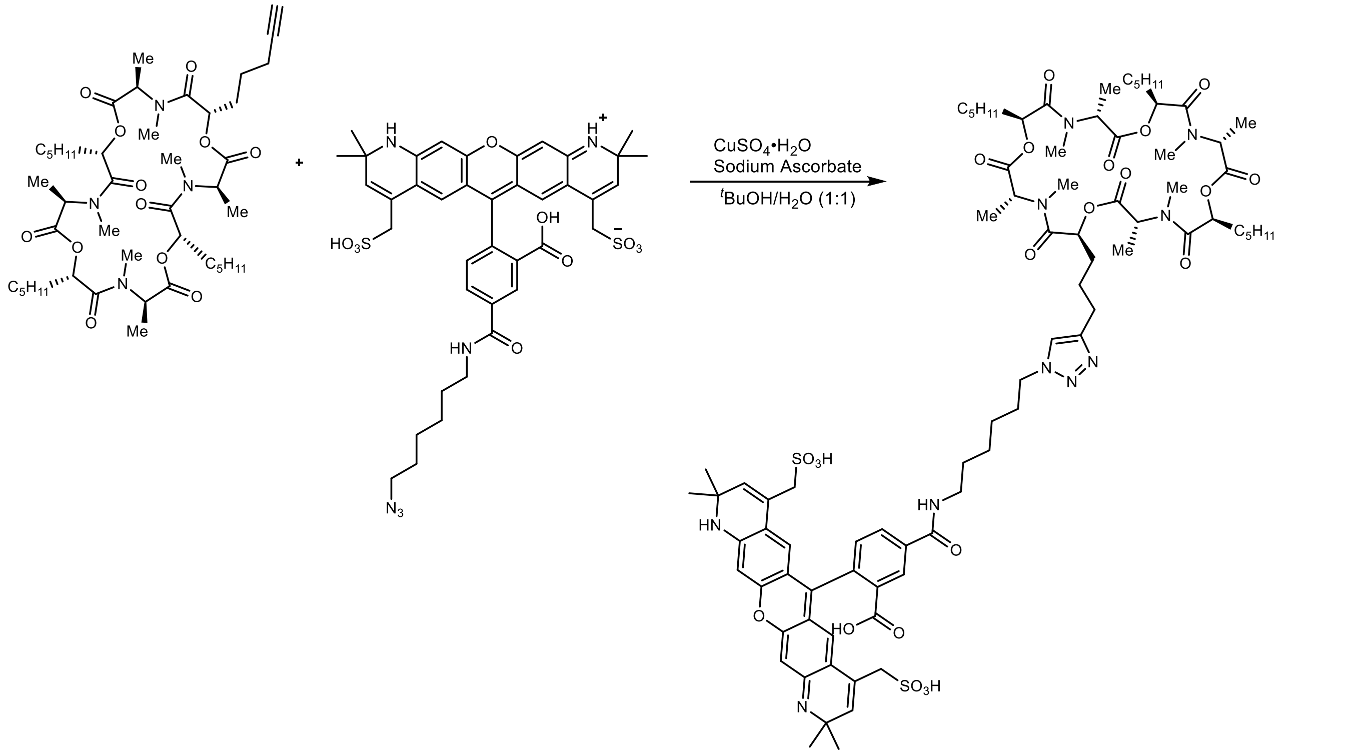

**5-((6-(4-(3-((2*S*,5*R*,8*S*,11*R*,14*S*,17*R*,20*S*,23*R*)-4,5,10,11,16,17,22,23-Octamethyl-3,6,9,12,15,18,21,24-octaoxo-8,14,20-tripentyl-1,7,13,19-tetraoxa-4,10,16,22-tetraazacyclotetracosan-2-yl)propyl)-1H-1,2,3-triazol-1-yl)hexyl)carbamoyl)-2-(2,2,10,10-tetramethyl-4,8-bis(sulfomethyl)-1,10-dihydro-2H-pyrano[3,2-g:5,6-g']diquinolin-6-yl)benzoic acid (3)**. A microwave vial was charged with the cyclic depsipeptide (8.9 mg, 11 µmol), azide (5.0 mg, 6.1 µmol), $^{t}$BuOH (500 µL), and $H_2O$ (500 µL). $CuSO_4{\cdot}H_2O$ (1 mg, 3.2 µmol) and sodium ascorbate (1.3 mg, 6.3 µmol) were added, the reaction was sealed with a microwave cap, and then submerged into a 50 °C oil bath. The reaction was allowed to stir at 50 °C for 2.5 h. The crude mixture was then cooled to ambient temperature, poured into $H_2O$, extracted with DCM (seven times). The combined organic layers were washed with 0.1 N aq EDTA·2Na and brine, and then dried and concentrated. Preparative HPLC (30 – 95% aqueous acetonitrile, 210 nm, flow rate: 8 mL/min, $R_t$ = 12.0 m) afforded the tagged 24-membered macrocycle after lyophilization as a bright purple amorphous solid (5.9 mg, 57%). A full characterization was not acquired due to limited material. However, a $^{1}$H NMR spectra is provided along with HRMS data; $^{1}$H NMR (600 MHz, $CH_3OD$) This compound exists in multiple conformations, causing significant peak overlap. Refer to the image of the $^{1}$H NMR spectrum for the other peaks: HRMS (EI): Exact mass calcd for $C_{83}H_{113}N_{10}O_{22}S_2$ [M-H]$^{-}$ 1665.7478, found 1665.7495. ANS-5-61.

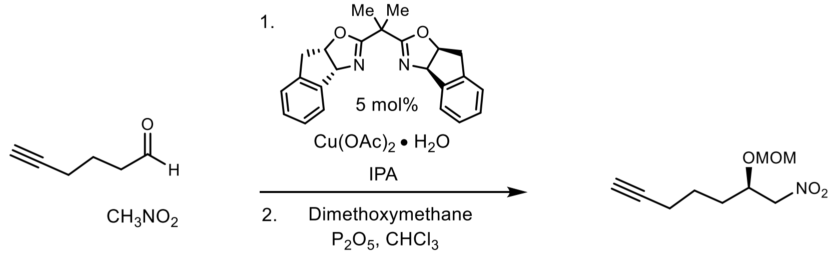

**(*R*)-6-(Methoxymethoxy)-7-nitrohept-1-yne ((*R*)-6).** Following the Evans enantioselective Henry procedure,[2] (1*R*,2*S*)-IndaBOX[3] (82.0 mg, 230 µmol) and $Cu(OAc)_2{\bullet}H_2O$ (41.5 mg, 210 µmol) were stirred at ambient temperature in isopropanol (8.3 mL) for 1 h. The cerulean blue solution was then cooled to 0 °C and heptanal (400 mg, 4.16 mmol) was added and

[2] Evans, D. A.; Seidel, D.; Rueping, M.; Lam, H. W.; Shaw, J. T.; Downey, C. W. *J. Am. Chem. Soc.* **2003**, *125*, 12692.
[3] Kurosu, M.; Porter, J. R.; Foley, M. A. *Tetrahedron Lett.* **2004**, *45*, 145.

allowed to stir for 10 m before nitromethane (2.20 mL, 41.6 mmol) addition. After stirring for 5 days at ambient temperature, the reaction was quenched dropwise at 0 °C with 1 M aq HCl and the aqueous layer was extracted with $CH_2Cl_2$. Following drying and concentration under reduced pressure, the crude alcohol was dissolved in $CHCl_3$ (20.8 mL), treated with $P_2O_5$ (5.90 g, 41.6 mmol) and dimethoxymethane (8.6 mL, 83.2 mmol), and stirred at ambient temperature overnight. The reaction mixture was diluted with DCM and decanted from the solid. The organic layer was then washed with satd aq $NaHCO_3$ and brine. The organic layers were dried and concentrated to afford an oil that was subjected to flash column chromatography ($SiO_2$, 3-6% diethyl ether in hexanes) to afford the title compound as a colorless oil (238 mg, 30%, 2 steps). The enantiopurity was determined to be 93% ee by chiral HPLC analysis (Chiralcel OD-H, 20% $^i$PrOH /hexanes, 0.4 mL/min, $t_r$($e_1$, major) = 15.6 min, $t_r$($e_2$, minor) = 17.6 min). $[\alpha]_D^{24}$ -14 (*c* 0.69, $CHCl_3$); $R_f$ = 0.19 (15% $Et_2O$/hexanes); IR (film) 3293, 2946, 1556, 1433, 1383, 1234, 1151, 1105, 1031 $cm^{-1}$; $^1$H NMR (400 MHz, $CDCl_3$) δ 4.68 (d, *J* = 7.2 Hz, 1H), 4.66 (d, *J* = 7.2 Hz, 1H), 4.52 (dd, *J*= 12.4, 8.0 Hz, 1H), 4.42 (dd, *J* = 12.4, 3.9 Hz, 1H), 4.34-4.27 (m, 1H), 3.36 (s, 3H), 2.25 (td, *J* = 6.8, 2.6 Hz, 2H), 1.98 (t, *J* = 2.7 Hz, 1H), 1.80-1.70 (m, 2H), 1.70-1.56 (m, 2H); $^{13}$C NMR (100 MHz, $CDCl_3$) ppm 96.5, 83.5, 79.1, 74.6, 69.3, 56.1, 31.5, 23.9, 18.4; HRMS (EI): Exact mass calcd for $C_9H_{14}NO_4$ $[M-H]^+$ 200.0917, found 200.0917. ANS-2-259.

**(*S*)-6-(Methoxymethoxy)-7-nitrohept-1-yne ((*S*)-6).** Prepared following an identical procedure as (*R*)-**6**, except using (1*S*,2*R*)-IndaBOX. Flash column chromatography ($SiO_2$, 3-6% diethyl ether in hexanes) afforded the hydroxy-nitroalkane with spectroscopic data identical to its enantiomer, except the major/minor peaks were reversed by chiral HPLC analysis. The enantiopurity was determined to be 92% ee by chiral HPLC analysis (Chiralcel OD-H, 20% $^i$PrOH /hexanes, 0.4 mL/min, $t_r$($e_1$, minor) = 15.6 min, $t_r$($e_2$, major) = 17.5 min). ANS-3-189.

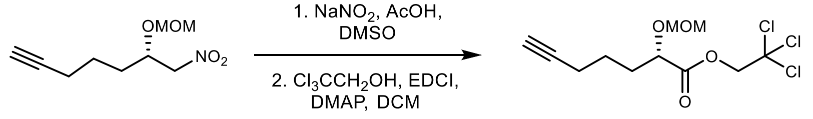

**2,2,2-Trichloroethyl (*S*)-2-(methoxymethoxy)hept-6-ynoate (8).** A round-bottom flask was charged with nitroalkane (823 mg, 4.09 mmol), $NaNO_2$ (847 mg, 12.3 mmol), AcOH (3.2 mL, 61.4 mmol), and DMSO (32 mL). The mixture was stirred at 60 °C for 18 h under an argon atmosphere. The reaction was then allowed to cool to ambient temperature, treated with 1 N HCl, and poured into $CH_2Cl_2$. The aqueous layer was extracted with $CH_2Cl_2$, and the combined organic layers were washed with water and brine, dried, and concentrated. The crude mixture was then added to a round-bottom flask and charged with trichloroethanol (376 µL, 3.90 mmol). The reaction was cooled to 0 °C and then treated with EDCI (855 mg, 4.46 mmol) and DMAP (13.6 mg, 112 µmol). The reaction was allowed to stir at 0 °C for 30 min, then warmed to ambient temperature and stirred for an additional 1.5 h. The reaction was poured into $H_2O$ and extracted with EtOAc. The organic layer was washed with water and brine, dried and concentrated. The crude residue was subjected to flash column chromatography ($SiO_2$, 10% ethyl acetate in hexanes) to afford the product as a pale-yellow oil (257 mg, 36%, 2 steps). $[\alpha]_D^{24}$ -52 (*c* 0.43, CHCl3); $R_f$ = 0.42 (20% $Et_2O$/hexanes); IR (film) 3301, 2953, 2898, 1765, 1443, 1150, 1114, 1032 $cm^{-1}$; $^1$H NMR (600 MHz, $CDCl_3$) δ 4.89 (d, *J* = 12.0 Hz, 1H), 4.74 (d, *J* = 7.1 Hz, 1H), 4.72 (d, *J* = 11.9 Hz, 1H), 4.70 (d, *J* = 7.0 Hz, 1H), 4.30 (dd, *J* = 7.9, 4.8 Hz, 1H), 3.41 (s, 3H), 2.26 (td, *J* = 7.0, 2.6 Hz, 2H), 2.05-1.91 (m, 2H), 1.96 (t, *J* = 2.6 Hz, 1H) 1.78-1.67 (m, 2H); $^{13}$C NMR (125 MHz, $CDCl_3$) ppm 171.1, 96.4, 94.8, 83.6, 74.8, 74.2, 69.1, 56.4, 31.8, 24.2, 18.2; HRMS (EI): Exact mass calcd for $C_{11}H_{16}Cl_3O_4$ $[M+H]^+$ 317.0109, found 307.0109. ANS-3-242.

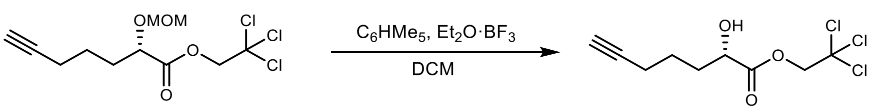

**2,2,2-Trichloroethyl (*S*)-2-hydroxyhept-6-ynoate (9).** A flame-dried round-bottom flask was charged with the protected alcohol (72.0 mg, 22.7 µmol) dissolved in dry dichloromethane (4.5 mL), pentamethyl benzene (67.3 mg, 45.4 µmol), and $BF_3 \cdot Et_2O$ (56 µL, 45 µmol). The reaction was allowed to stir for 1 h and 20 min at ambient temperature. The crude reaction mixture was quenched with satd aq $NaHCO_3$, washed with brine, dried, concentrated, and subjected to flash column chromatography ($SiO_2$, 20% ethyl acetate in hexanes) to afford the alcohol (23.4 mg, 38%) as a pale-yellow oil. $[\alpha]_D^{23}$ -9.15 (*c* 1.05, $CHCl_3$); $R_f$ = 0.17 (20% EtOAc/hexanes); IR (film) 3507, 3303, 2954, 1756, 1582, 1439, 1269, 1167, 1109 $cm^{-1}$; $^1H$ NMR (400 MHz, $CDCl_3$) δ 4.93 (d, *J* = 11.8 Hz, 1H), 4.73 (d, *J* = 11.8 Hz, 1H), 4.38 (ddd, *J* = 7.4, 5.7, 4.4 Hz, 1H), 2.65 (d, *J* = 6.0 Hz, 1H), 2.27 (td, *J* = 6.9, 2.6 Hz, 2H), 2.10-1.99 (m, 1H), 1.96 (t, *J* = 2.6 Hz, 1H) 1.91-1.64 (series of m, 3H); $^{13}C$ NMR (125 MHz, $CDCl_3$) ppm 173.7, 94.5, 83.7, 74.6, 70.2, 69.1, 33.3, 23.9, 18.2; HRMS (EI): Exact mass calcd for $C_9H_{12}Cl_3O_3$ $[M+H]^+$ 272.9847, found 272.9847. ANS-3-234.

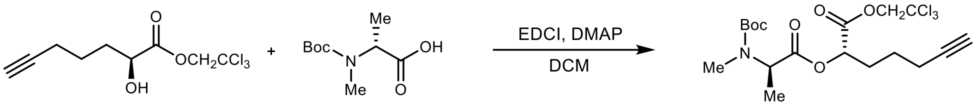

**2,2,2-Trichloroethyl (*S*)-2-((*N*-(*tert*-butoxycarbonyl)-*N*-methyl-D-alanyl)oxy)hept-6-ynoate (10).** A round-bottom flask was charged with the amine (20 mg, 98 µmol), alcohol (23.4 mg, 85.6 µmol), and DCM (1.0 mL). The mixture was cooled to 0 °C and then EDCI (19.7 mg, 103 µmol) and DMAP (1.0 mg, 4.3 µmol) were added. The reaction was stirred at 0 °C for 30 min, then allowed to warm to ambient temperature and stir for an additional 1.5 h. The reaction mixture was poured into water and extracted with DCM. The combined organic layers were washed with satd aq $NaHCO_3$ and brine, dried, and concentrated to afford the product as a colorless oil (34.9 mg, 88%). $[\alpha]_D^{24}$ +5.0 (*c* 0.87, $CHCl_3$); $R_f$ = 0.39 (20% EtOAc/hexanes); IR (film) 2973, 1750, 1695, 1452, 1390, 1156, 1089 $cm^{-1}$; $^1H$ NMR (600 MHz, $CDCl_3$) This compound exists as a 1.4:1 ratio of rotamers causing significant peak broadening and overlap. Refer to the image of the $^1H$ NMR spectrum. $^{13}C$ NMR (150 MHz, $CDCl_3$) This compound is a 1.4:1 ratio of rotamers causing significant peak broadening and overlap. Refer to the image of the $^{13}C$ NMR spectrum; HRMS (EI): Exact mass calcd for $C_{18}H_{26}Cl_3NO_6Na$ $[M+Na]^+$ 480.0718, found 480.0731. ANS-3-243.

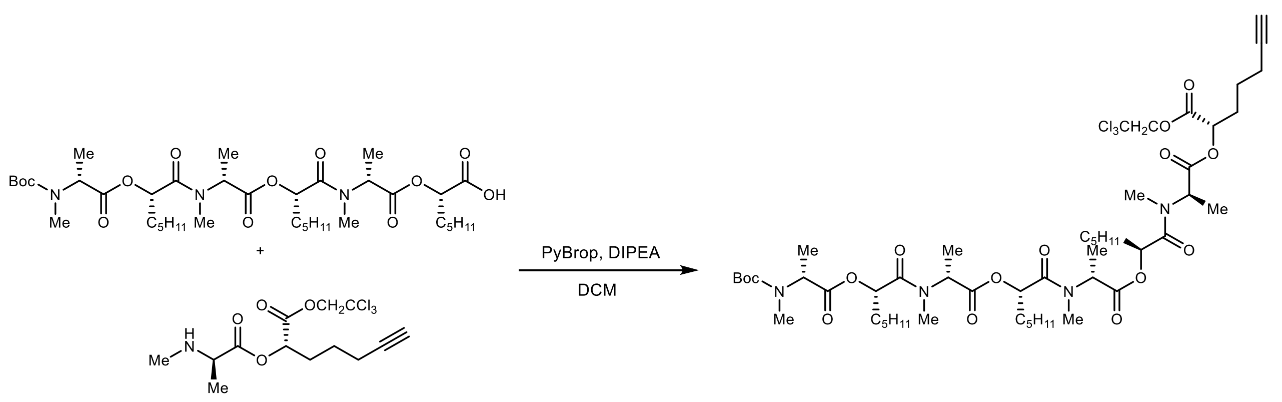

**2,2,2-Trichloroethyl (*S*)-2-(((6*R*,9*S*,12*R*,15*S*,18*R*,21*S*,24*R*)-2,2,5,6,11,12,17,18,23,24-decamethyl-4,7,10,13,16,19,22-heptaoxo-9,15,21-tripentyl-3,8,14,20-tetraoxa-5,11,17,23-tetraazapentacosan-25-oyl)oxy)hept-6-ynoate (12)**. A round-bottom flask was charged with the amine (14.8 mg, 41.5 µmol), acid (33.9 mg, 44.8 µmol), and DCM (450 µL). The mixture was cooled to 0 °C and then DIPEA (23.0 µL, 134 µmol) and PyBrop (31.3 mg, 67.2 µmol) were added. The reaction was stirred at 0 °C for 30 min, then allowed to warm to ambient temperature and stir for an additional 1.5 h. The reaction mixture was poured into cold 10% aq citric acid and extracted with DCM. The combined organic layers were washed with satd aq $NaHCO_3$ and brine, dried, and concentrated. The crude residue was subjected to flash column chromatography ($SiO_2$, 30-50% ethyl acetate in hexanes) to afford the product as a colorless oil (33.2 mg, 67%). $[\alpha]_D^{24}$ +38 (*c* 0.48, $CHCl_3$); $R_f$ = 0.21 (40% EtOAc/hexanes); IR (film) 2930, 2864, 1742, 1663, 1457, 1384, 1216, 1159, 1085 $cm^{-1}$; $^1H$ NMR (600 MHz, $CDCl_3$) This compound exists as a mixture of rotamers causing significant peak broadening and overlap. Refer to the image of the $^1H$ NMR spectrum; $^{13}C$ NMR (150 MHz, $CDCl_3$) This compound is a mixture of rotamers causing significant peak broadening and overlap. Refer to the image of the $^{13}C$ NMR spectrum; HRMS (EI): Exact mass calcd for $C_{51}H_{83}Cl_3N_4NaO_{15}$ $[M+Na]^+$ 1119.4813, found 1119.4836. ANS-3-263.

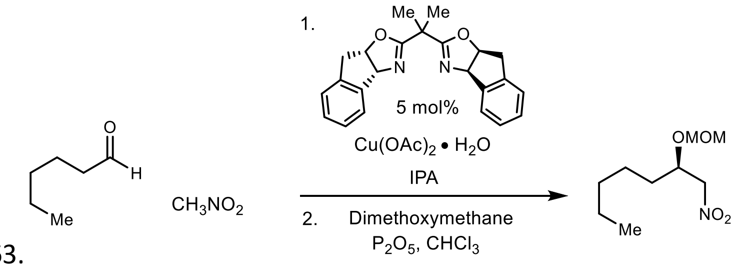

**(*R*)-2-(methoxymethoxy)-1-nitroheptane ((*R*)-S1).** Following the Evans protocol,[2] (1*R*,2*S*)-IndaBOX[3] (321 mg, 897 µmol) and $Cu(OAc)_2$•$H_2O$ (162 mg, 813 µmol) were stirred at ambient temperature in isopropanol (32.6 mL) for 1 h. The cerulean blue solution was then cooled to 0 °C, and hexanal (2.00 mL, 16.3 mmol) was added and allowed to stir for 10 m before nitromethane (9.95 mL, 163 mmol) addition. After stirring for 4 days at ambient temperature, the reaction was quenched dropwise at 0 °C with 1 M aq HCl and the aqueous layer was extracted with $CH_2Cl_2$. Following drying and concentration under reduced pressure, the crude alcohol was dissolved in $CHCl_3$ (81.6 mL), treated with $P_2O_5$ (23.1 g, 163 mmol) and dimethoxymethane (33.9 mL, 326 mmol), and stirred at ambient temperature overnight. The reaction mixture was diluted with DCM and decanted from the solid. The organic layer was then washed with satd aq $NaHCO_3$ and brine. The organic layers were dried and concentrated to afford an oil that was subjected to flash column chromatography ($SiO_2$, 3-6% diethyl ether in hexanes) to afford the title compound as a pale-yellow oil (2.54 g, 76%, 2 steps). All spectral data are in agreement with literature values.[4] ANS-2-254.

**(*S*)-2-(methoxymethoxy)-1-nitroheptane ((*S*)-S1).** Prepared following an identical procedure as **1**, except using (1*S*,2*R*)-IndaBOX. Flash column chromatography ($SiO_2$, 3-6% ethyl acetate in hexanes) afforded the hydroxy-nitroalkane with spectroscopic data identical to its enantiomer, except the major/minor peaks were reversed by chiral HPLC analysis. The enantiopurity was determined to be 95% ee by chiral HPLC analysis (Chiralcel OD-H, 2% $^i$PrOH /hexanes, 0.4 mL/min, $t_r(e_1$, minor) = 16.7 min, $t_r(e_2$, major) = 19.1 min). ANS-2-291.

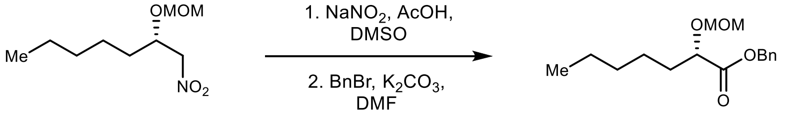

**Benzyl (*S*)-2-(methoxymethoxy)heptanoate (S2).** A round-bottom flask was charged with (*S*)-nitroalkane (1.36 g, 6.63 mmol), $NaNO_2$ (1.37 g, 19.9 mmol), AcOH (5.7 mL, 99.5 mmol), and DMSO (51.0 mL). The mixture was stirred at 45 °C for 18 h under an argon atmosphere. The reaction was then allowed to cool to ambient temperature, treated with 1 N HCl, and poured into $CH_2Cl_2$. The aqueous layer was extracted with $CH_2Cl_2$, and the combined organic layers were washed with water, brine, dried, and concentrated. The crude mixture was then added to a round-bottom flask with $K_2CO_3$ (2.75 g, 19.9 mmol) and DMF (13.2 mL). The mixture was then treated with BnBr (950 µL, 7.95 mmol) and the reaction was allowed to stir overnight at ambient temperature under an argon atmosphere (balloon). The reaction was quenched with 1 M aq HCl and extracted with $Et_2O$. The organic layer was washed with 1 M HCl, water, brine, dried and concentrated. The crude residue was subjected to flash column chromatography ($SiO_2$, 5% diethyl ether in hexanes) to afford the product as a pale-yellow oil (1.30 g, 70%, 2 steps). $[\alpha]_D^{20}$ -52.2 (*c* 1.04, $CHCl_3$); $R_f$ = 0.22 (15% $Et_2O$/hexanes); IR (film) 2953, 2863, 1749, 1457, 1262, 1156, 1126, 1037 $cm^{-1}$; $^1H$ NMR (400 MHz, $CDCl_3$) δ 7.40-7.26 (m, 5H), 5.19 (d, *J* = 12.4 Hz, 1H), 5.15 (d, *J*= 12.3 Hz, 1H), 4.68 (d, *J* = 6.9 Hz, 1H), 4.65 (d, *J* = 6.9 Hz, 1H), 4.14 (dd, *J* = 6.3, 6.3 Hz, 1H), 3.34 (s, 3H), 1.75 (dddd, *J* = 7.3, 7.3, 7.3, 7.3 Hz, 2H), 1.45-1.20 (m, 6H), 0.86 (t, *J* = 6.6 Hz, 3H); $^{13}C$ NMR (100 MHz, $CDCl_3$) ppm 172.7, 135.7, 128.6, 128.4, 128.3,

[4] Batiste, S. M.; Johnston, J. N. *Proc. Natl. Acad. Sci. U. S. A.* **2016**, *113*, 14893.

96.3, 75.7, 66.5, 56.0, 32.9, 31.5, 24.9, 22.5, 14.0; HRMS (EI): Exact mass calcd for $C_{16}H_{24}O_4Na$ $[M+Na]^+$ 303.1567, found 303.1569. ANS-2-273.

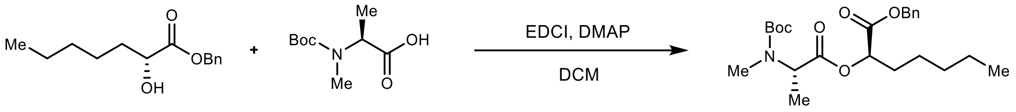

**Benzyl (*R*)-2-((*N*-(*tert*-butoxycarbonyl)-*N*-methyl-L-alanyl)oxy)heptanoate ((*R*)-S3).** A round-bottom flask was charged with the amine (1.19 g, 5.84 mmol), alcohol (1.20 g, 5.08 mmol), and DCM (51 mL). The mixture was cooled to 0 °C and then EDCI (1.17 g, 6.09 mmol) and DMAP (31.0 mg, 254 μmol) were added. The reaction was stirred at 0 °C for 30 min, then allowed to warm to ambient temperature and stir for an additional 1.5 h. The reaction mixture was poured into water and extracted with DCM. The combined organic layers were washed with satd aq $NaHCO_3$ and brine, and then dried and concentrated. The crude residue was subjected to flash column chromatography ($SiO_2$, 5% ethyl acetate in hexanes) to afford the product as a colorless oil (1.88 g, 88%). All spectral data are in agreement with literature values.[5] ANS-3-08.

**Benzyl (*S*)-2-((*N*-(*tert*-butoxycarbonyl)-*N*-methyl-D-alanyl)oxy)heptanoate ((*S*)-S3).** Prepared following an identical procedure as (*R*)-**S3**. Flash column chromatography ($SiO_2$, 5% ethyl acetate in hexanes) afforded the depsipeptide with spectroscopic data identical to its enantiomer, except $[\alpha]_D^{26}$ +12 (*c* 0.94, $CHCl_3$). ANS-2-282.

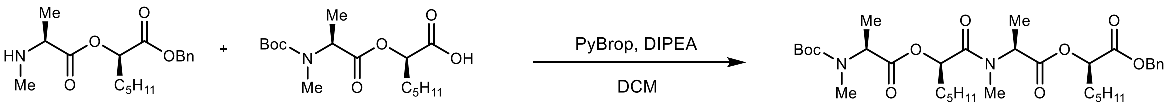

**Benzyl (*R*)-2-(((6*S*,9*R*,12*S*)-2,2,5,6,11,12-hexamethyl-4,7,10-trioxo-9-pentyl-3,8-dioxa-5,11-diazatridecan-13-oyl)oxy)heptanoate ((*R*)-S4).** A round-bottom flask was charged with the amine (231 mg, 717 μmol), acid (236 mg, 717 μmol), and DCM (4.0 mL). The mixture was cooled to 0 °C and then DIPEA (370 μL, 2.15 mmol) and PyBrop (502 mg, 1.08 mmol) were added. The reaction was stirred at 0 °C for 1 h. The reaction mixture was poured into cold 10% citric acid and extracted with DCM. The combined organic layers were washed with satd aq $NaHCO_3$ and brine, and then dried and concentrated. The crude residue was subjected to flash column chromatography ($SiO_2$, 10-40% ethyl acetate in hexanes) to afford the product as a colorless oil (330 mg, 73%). All spectral data are in agreement with literature values.[5] ANS-3-19.

**Benzyl (*S*)-2-(((6*R*,9*S*,12*R*)-2,2,5,6,11,12-hexamethyl-4,7,10-trioxo-9-pentyl-3,8-dioxa-5,11-diazatridecan-13-oyl)oxy)heptanoate ((*S*)-S4).** Prepared following an identical procedure as (*R*)-**S4**. Flash column chromatography ($SiO_2$, 10-40% ethyl acetate in hexanes) afforded the depsipeptide with spectroscopic data identical to its enantiomer, except $[\alpha]_D^{26}$ +28.0 (*c* 1.00, $CHCl_3$). ANS-2-288.

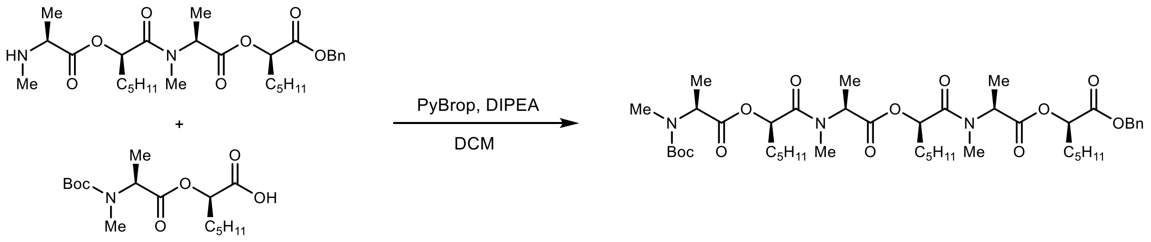

**Benzyl (*R*)-2-(((6*S*,9*R*,12*S*,15*R*,18*S*)-2,2,5,6,11,12,17,18-octamethyl-4,7,10,13,16-pentaoxo-9,15-dipentyl-3,8,14-trioxa-5,11,17-triazanonadecan-19-oyl)oxy)heptanoate ((*R*)-S5).** A round-bottom flask was charged with the amine (21.1 mg, 39.5 μmol), acid (13.1 mg, 39.5 μmol), and DCM (0.4 mL). The mixture was cooled to 0 °C and then DIPEA (20.3 μL, 119 μmol) and PyBrop (28.0 mg, 59.3 μmol) were added. The reaction was stirred at 0 °C for 30 min, then allowed to warm to

[5] Monma, et al., *Org. Lett.* **2006,** 5601.

ambient temperature and stir for an additional 1.5 h. The reaction mixture was poured into cold 10% citric acid and extracted with DCM. The combined organic layers were washed with satd aq $NaHCO_3$ and brine, and then dried and concentrated. The crude residue was subjected to flash column chromatography ($SiO_2$, 10-50% ethyl acetate in hexanes) to afford the product as a colorless oil (32 mg, 96%). $[\alpha]_D^{25}$ – 34 (*c* 0.95, $CHCl_3$); $R_f$ = 0.14 (20% EtOAc/hexanes); IR (film) 2932, 2865, 1743, 1666, 1458, 1316, 1187, 1086 $cm^{-1}$; $^1H$ NMR (600 MHz, $CDCl_3$) This compound exists as a mixture of rotamers causing significant peak broadening and overlap. Refer to image of the $^1H$ NMR spectrum; $^{13}C$ NMR (150 MHz, $CDCl_3$) This compound is a mixture of rotamers causing significant peak broadening and overlap. Refer to the image of the $^{13}C$ NMR spectrum; HRMS (EI): Exact mass calcd for $C_{45}H_{73}N_3NaO_{12}$ $[M+Na]^+$ 870.5092, found 870.5071. ANS-3-31.

**Benzyl (*S*)-2-(((6*R*,9*S*,12*R*,15*S*,18*R*)-2,2,5,6,11,12,17,18-octamethyl-4,7,10,13,16-pentaoxo-9,15-dipentyl-3,8,14-trioxa-5,11,17-triazanonadecan-19-oyl)oxy)heptanoate ((*S*)-S5).** Prepared following an identical procedure as (*R*)-**S5**. Flash column chromatography ($SiO_2$, 10-50% ethyl acetate in hexanes) afforded the protected depsipeptide with spectroscopic data identical to its enantiomer, except $[\alpha]_D^{20}$ +33 (*c* 0.90, $CHCl_3$). ANS-3-150.

**Figure S1. $^1$H NMR (600 MHz, $CDCl_3$) of 2**

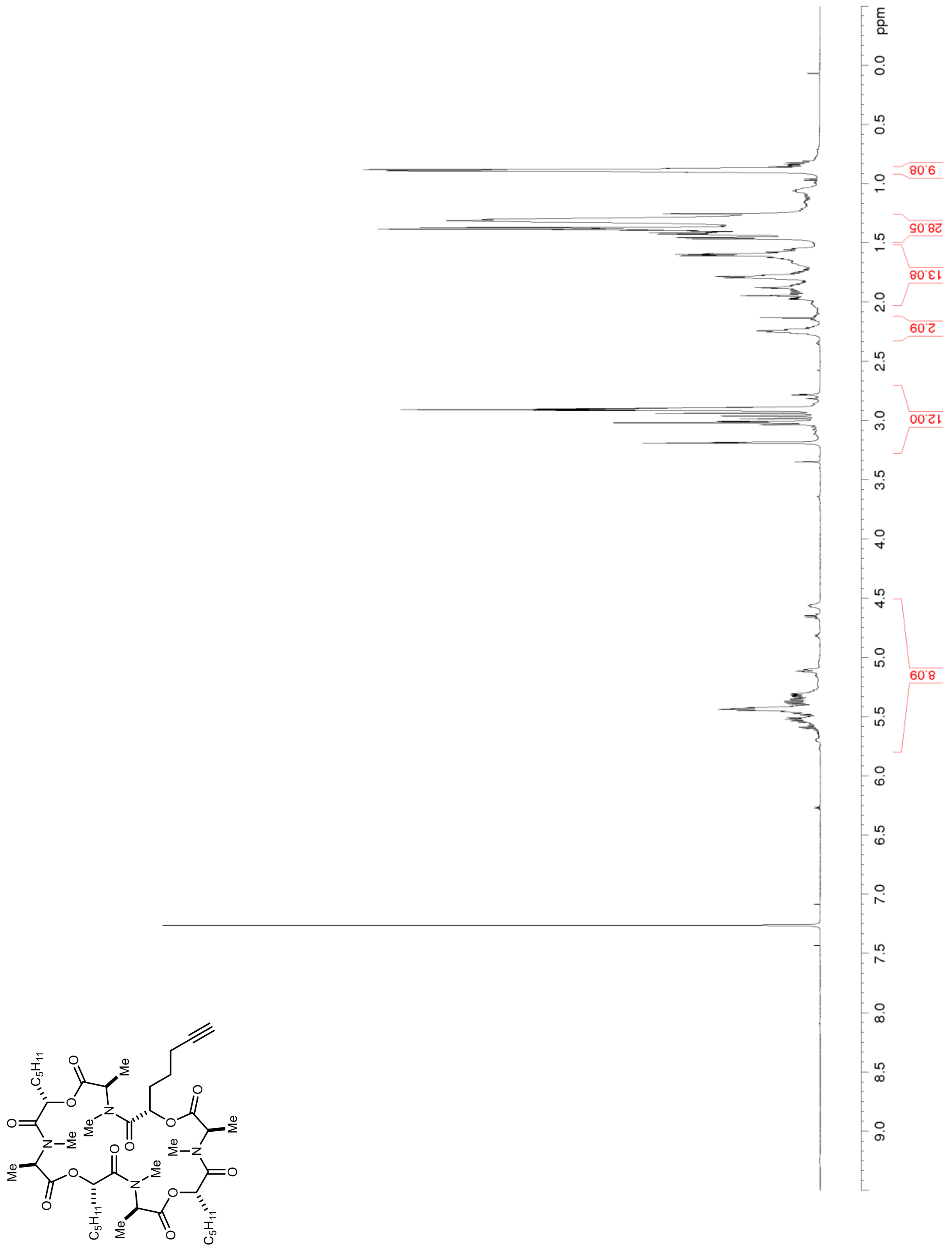

**Figure S2. $^{13}C$ NMR/DEPT (150 MHz, $CDCl_3$) of 2**

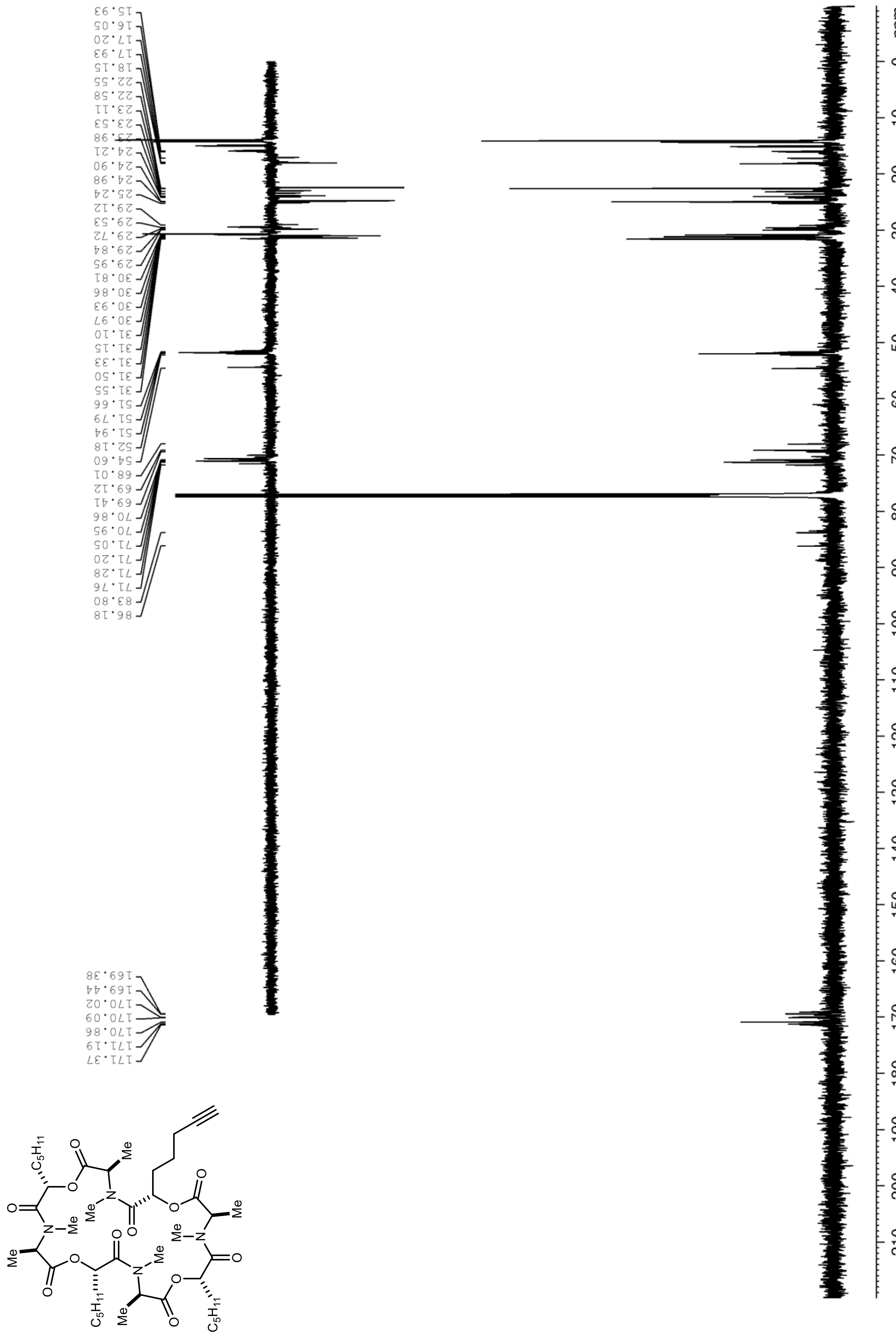

**Figure S3. $^1$H NMR (600 MHz, $CDCl_3$) of 3**

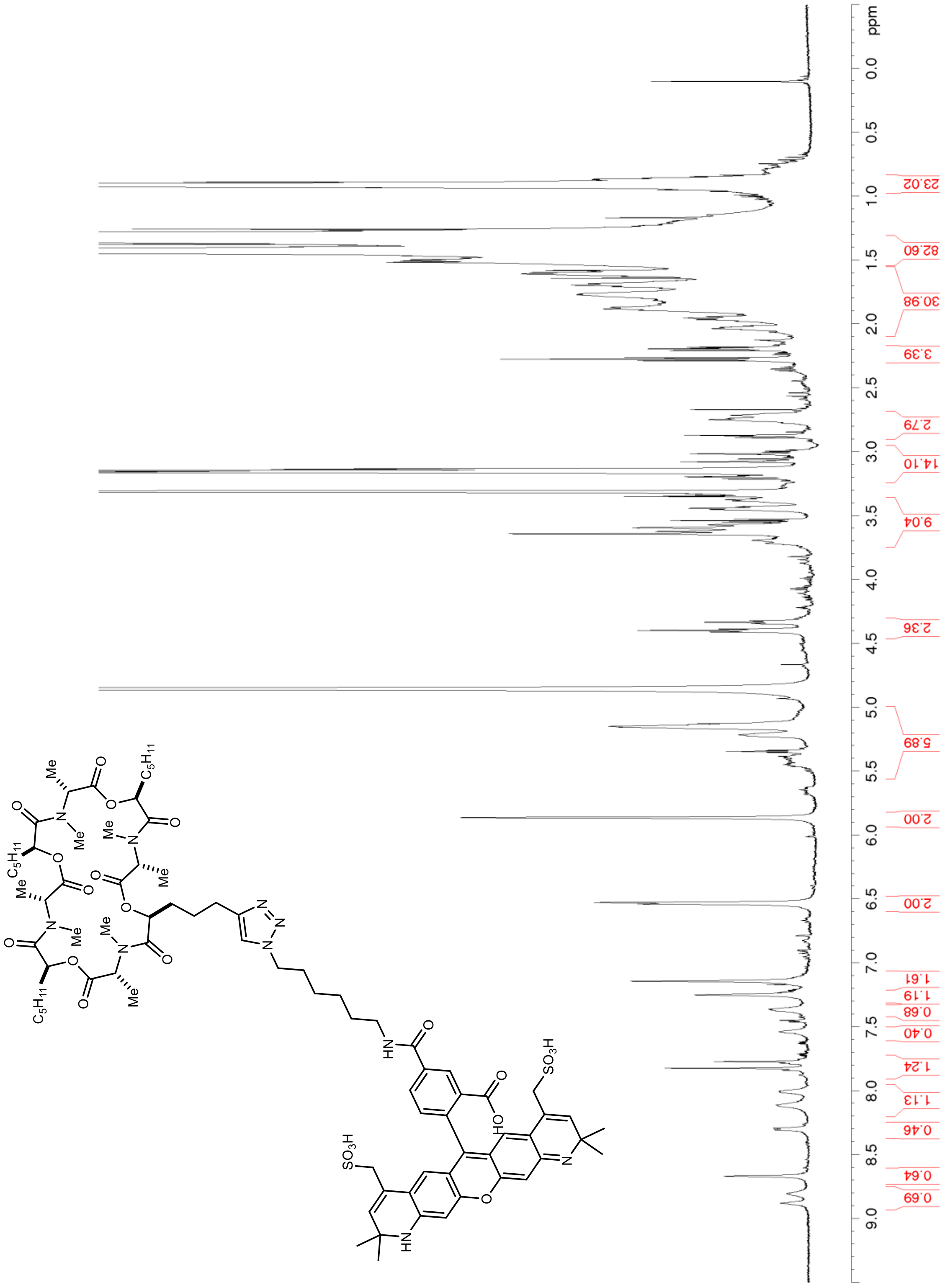

**Figure S4.** $^{1}$**H NMR (400 MHz,** $CDCl_3$**) of 6**

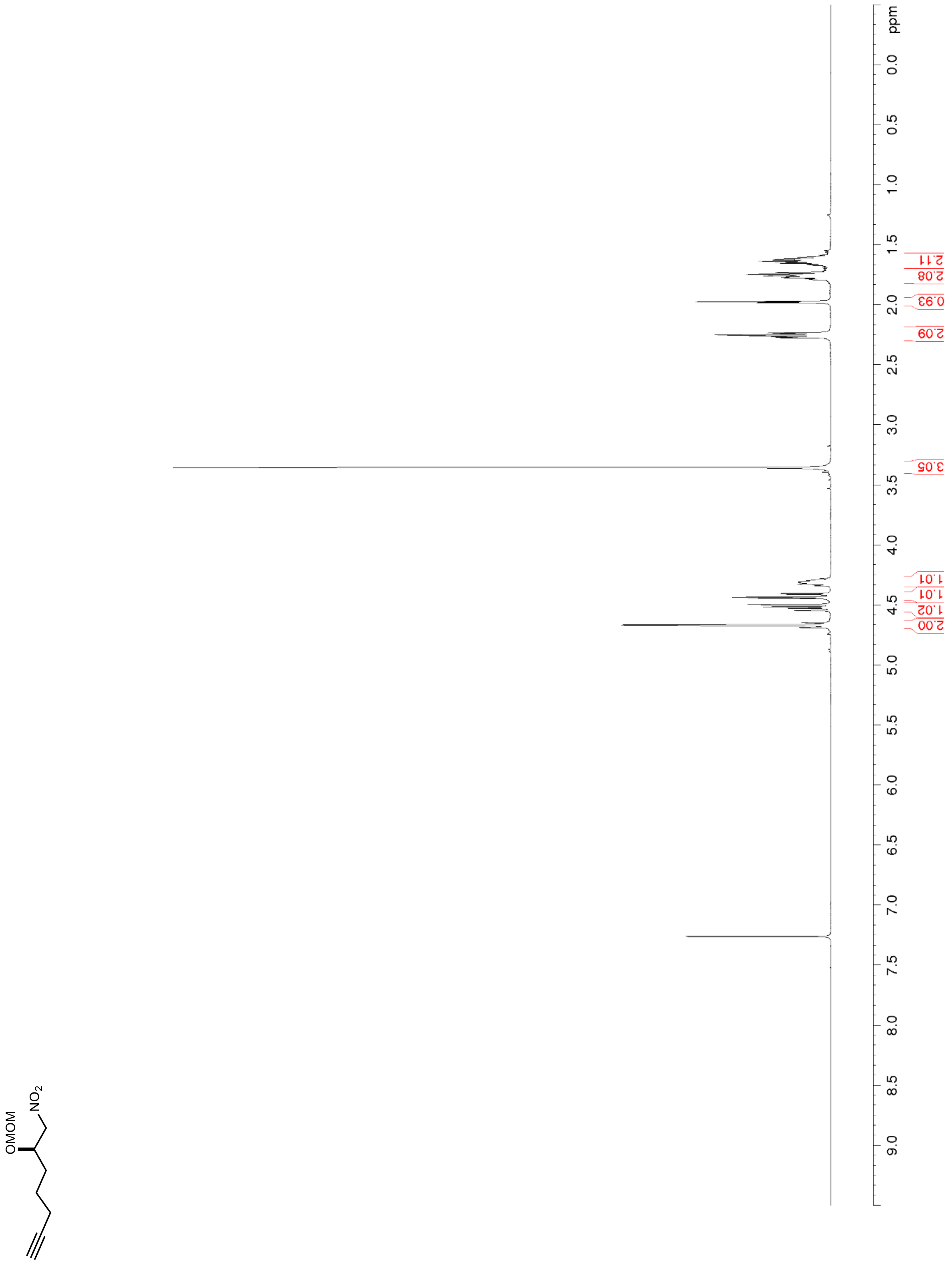

**Figure S5. $^{13}$C NMR/DEPT (100 MHz, $CDCl_3$) of 6**

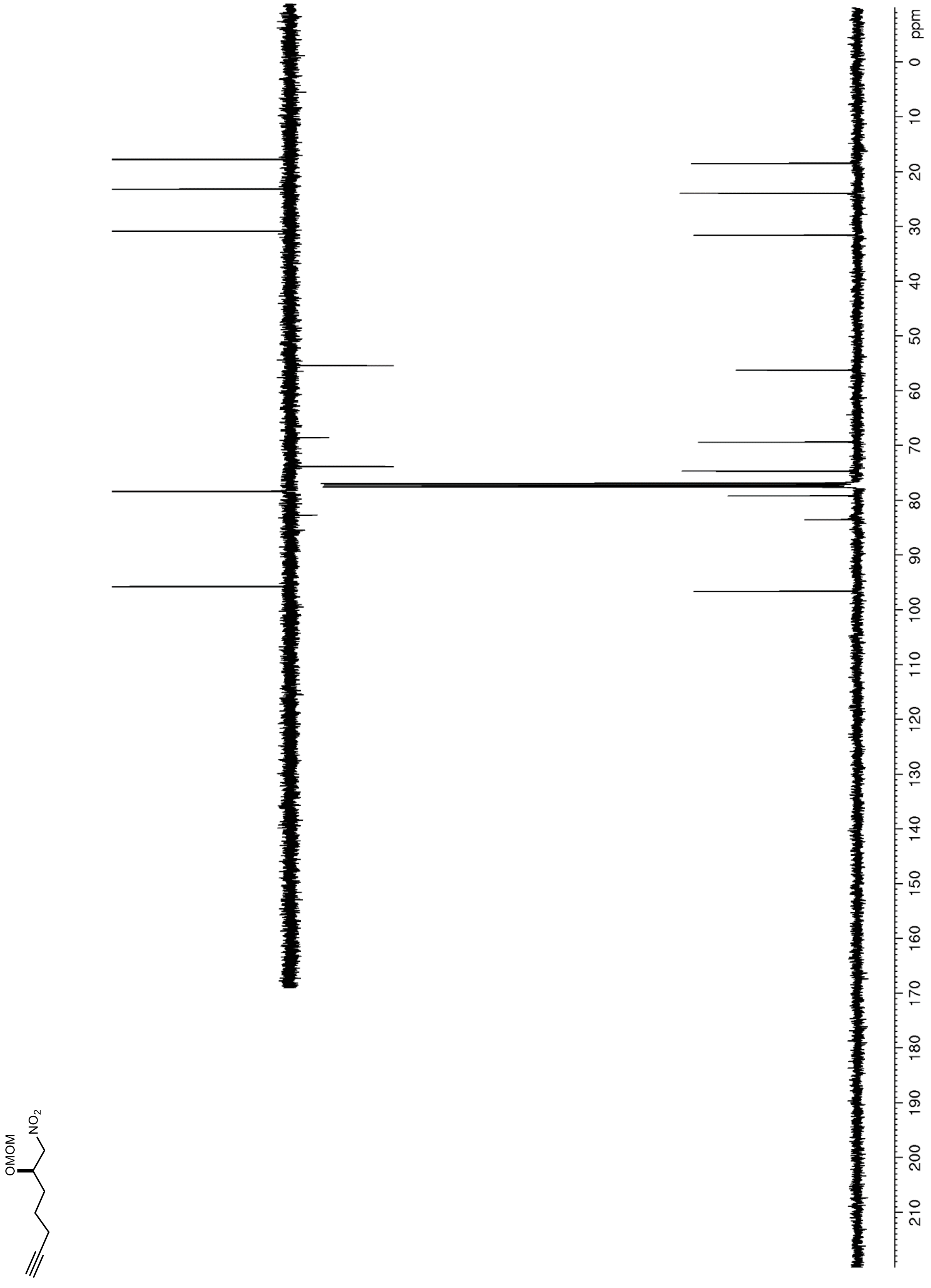

**Figure S6. $^{1}$H NMR (600 MHz, $CDCl_3$) of 8**

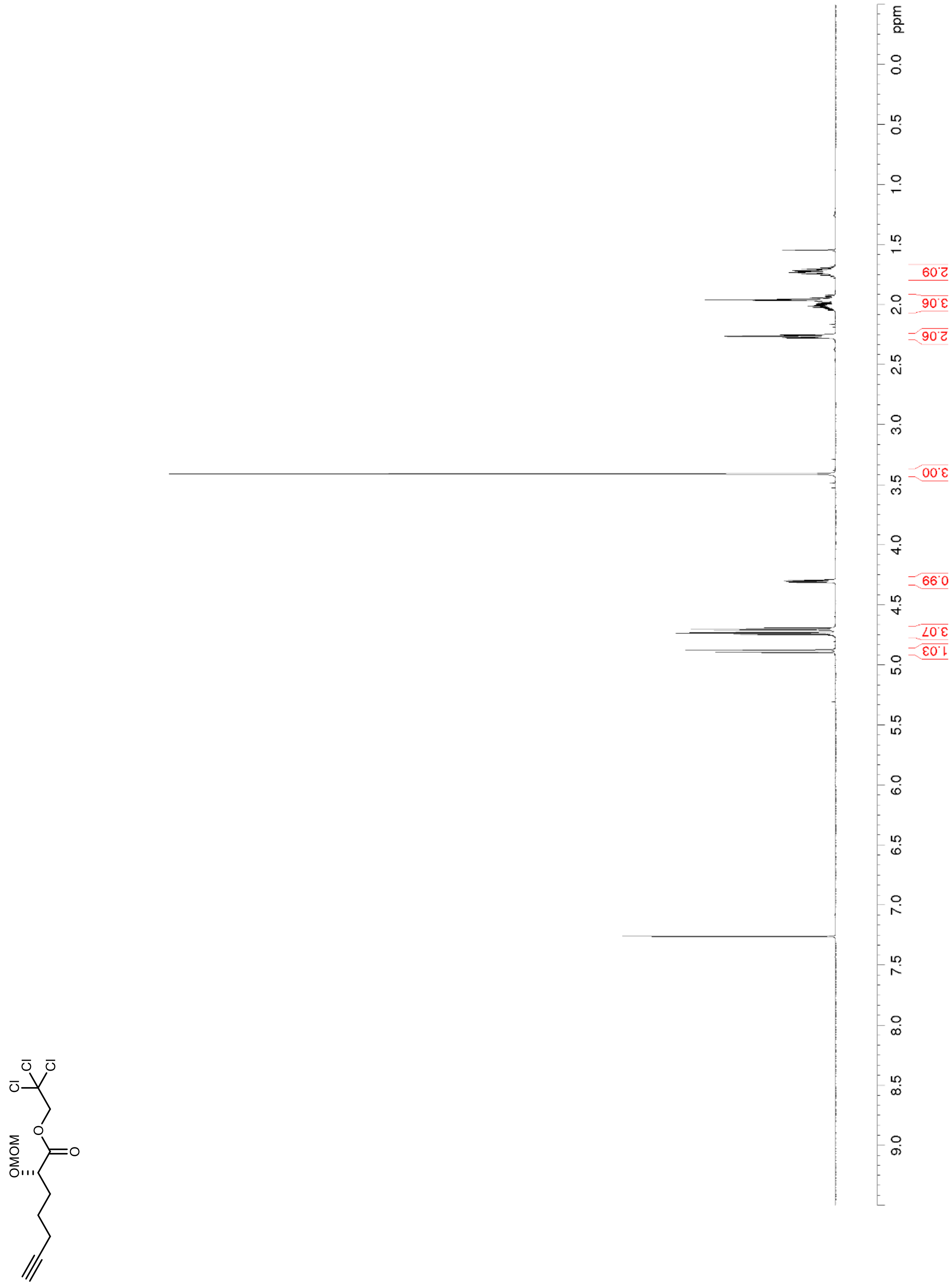

**Figure S7. $^{13}C$ NMR/DEPT (150 MHz, $CDCl_3$) of 8**

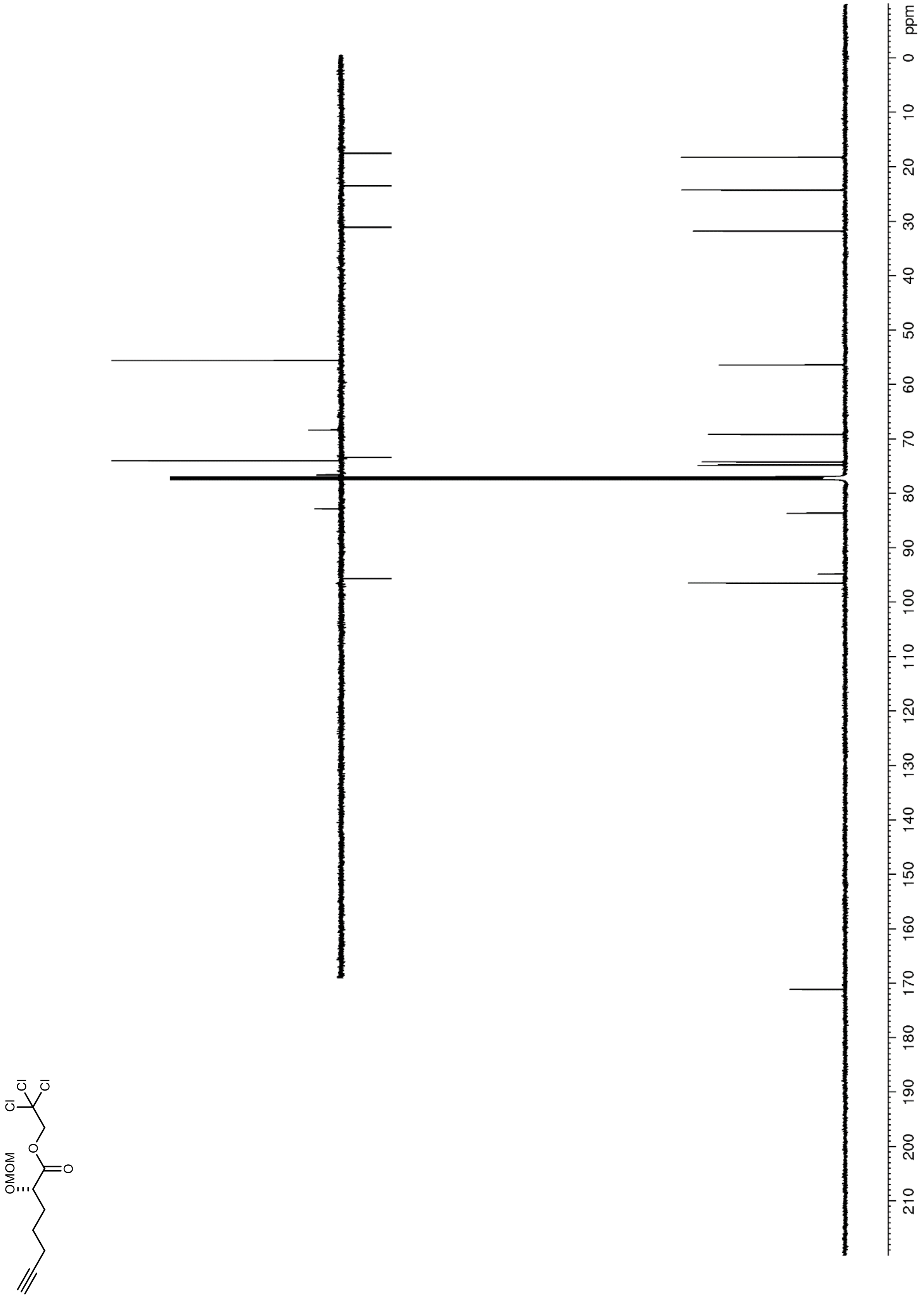

**Figure S8.** **$^{1}H$ NMR (400 MHz, $CDCl_3$) of 9**

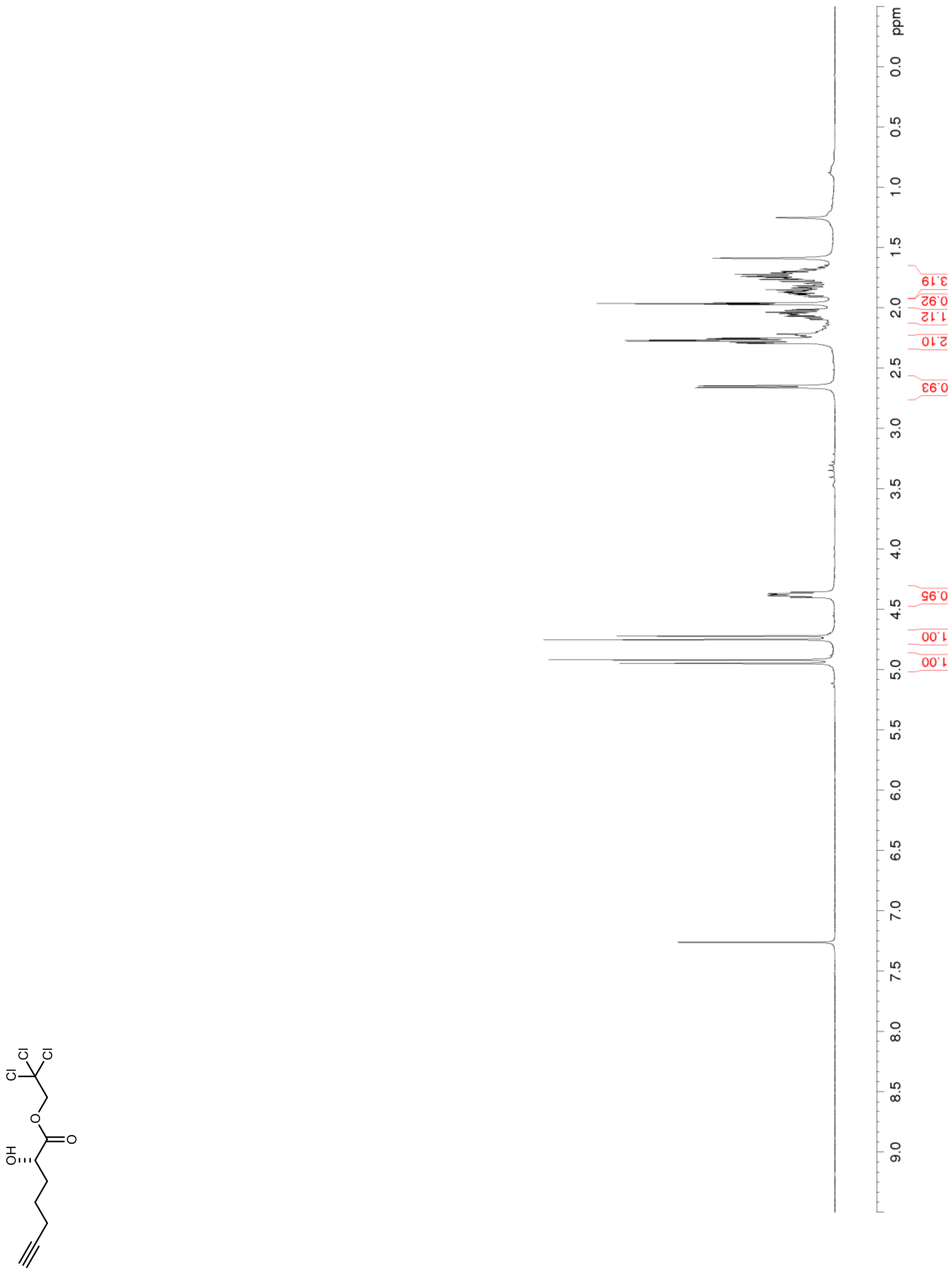

**Figure S9. $^{13}$C NMR/DEPT (100 MHz, $CDCl_3$) of 9**

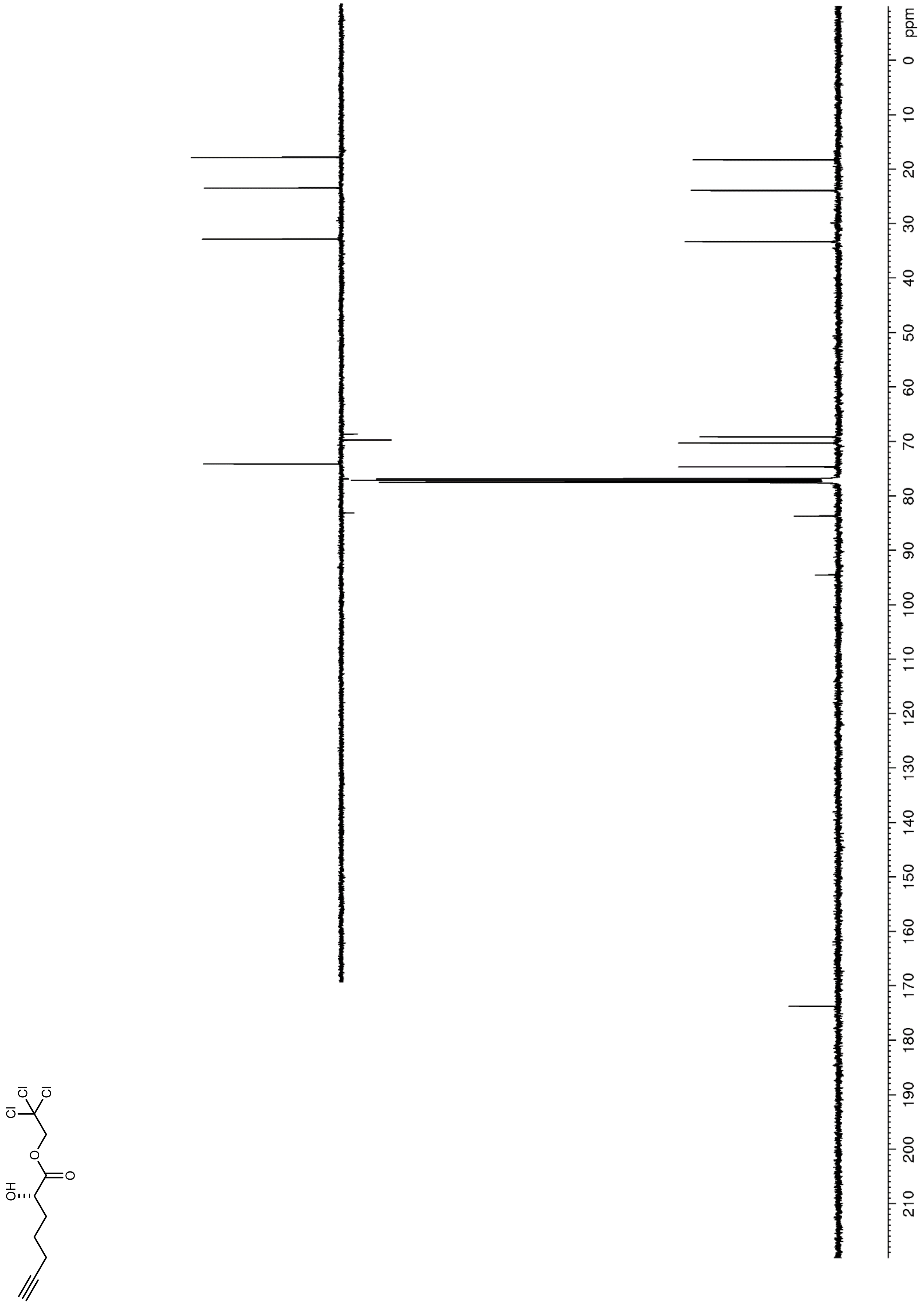

**Figure S10. $^1$H NMR (600 MHz, $CDCl_3$) of 10**

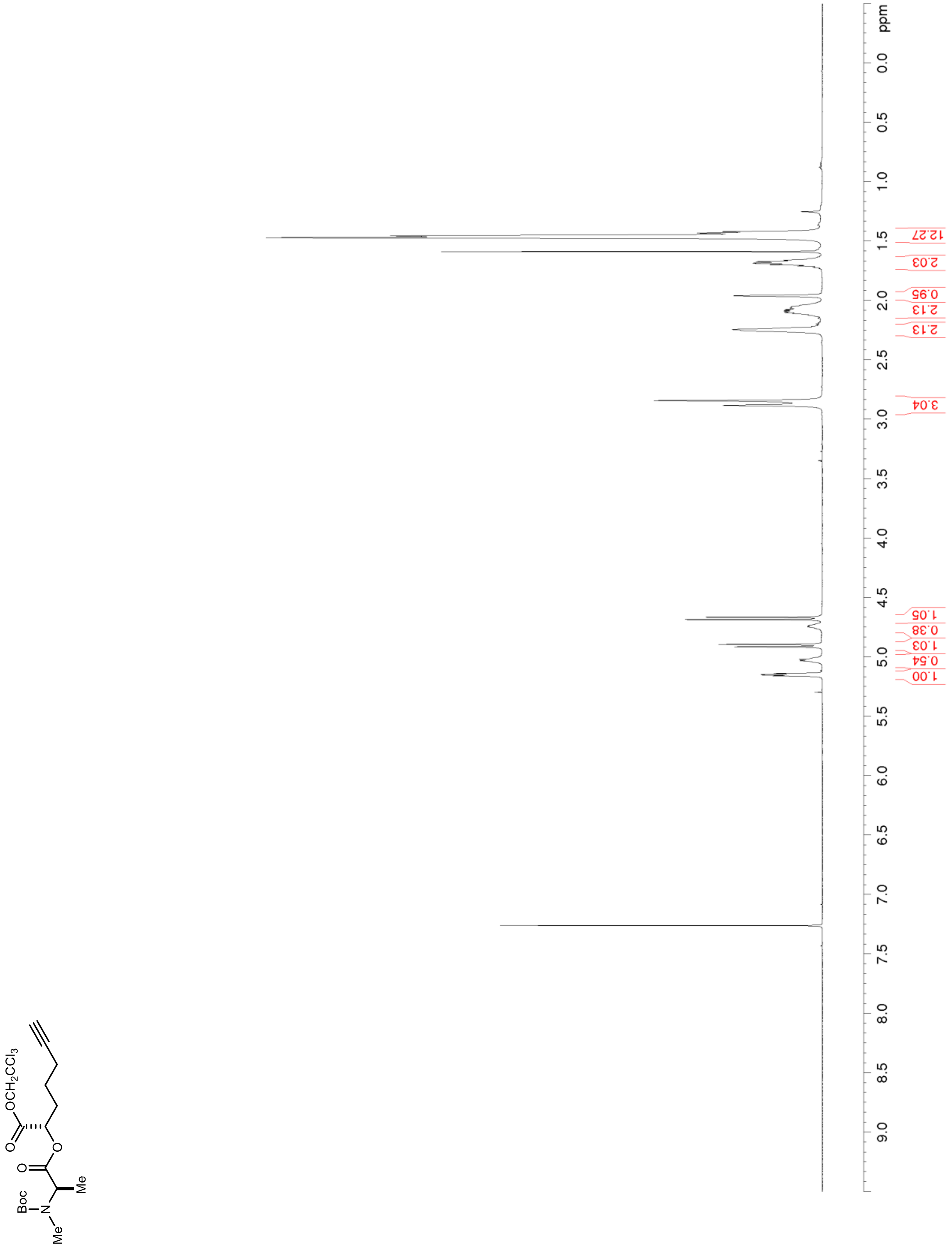

**Figure S11. $^{13}C$ NMR/DEPT (150 MHz, $CDCl_3$) of 10**

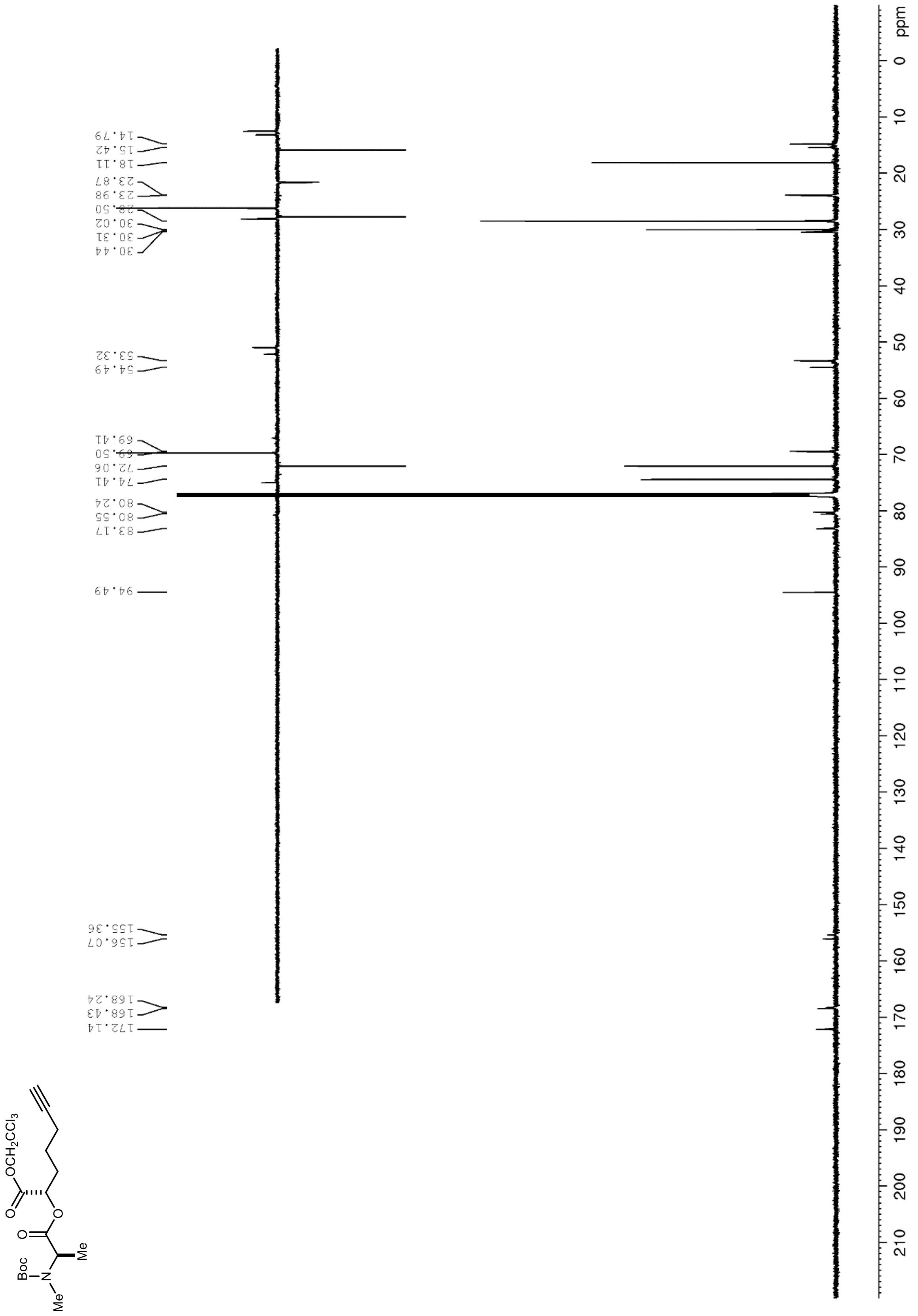

**Figure S12. $^1H$ NMR (600 MHz, $CDCl_3$) of 12**

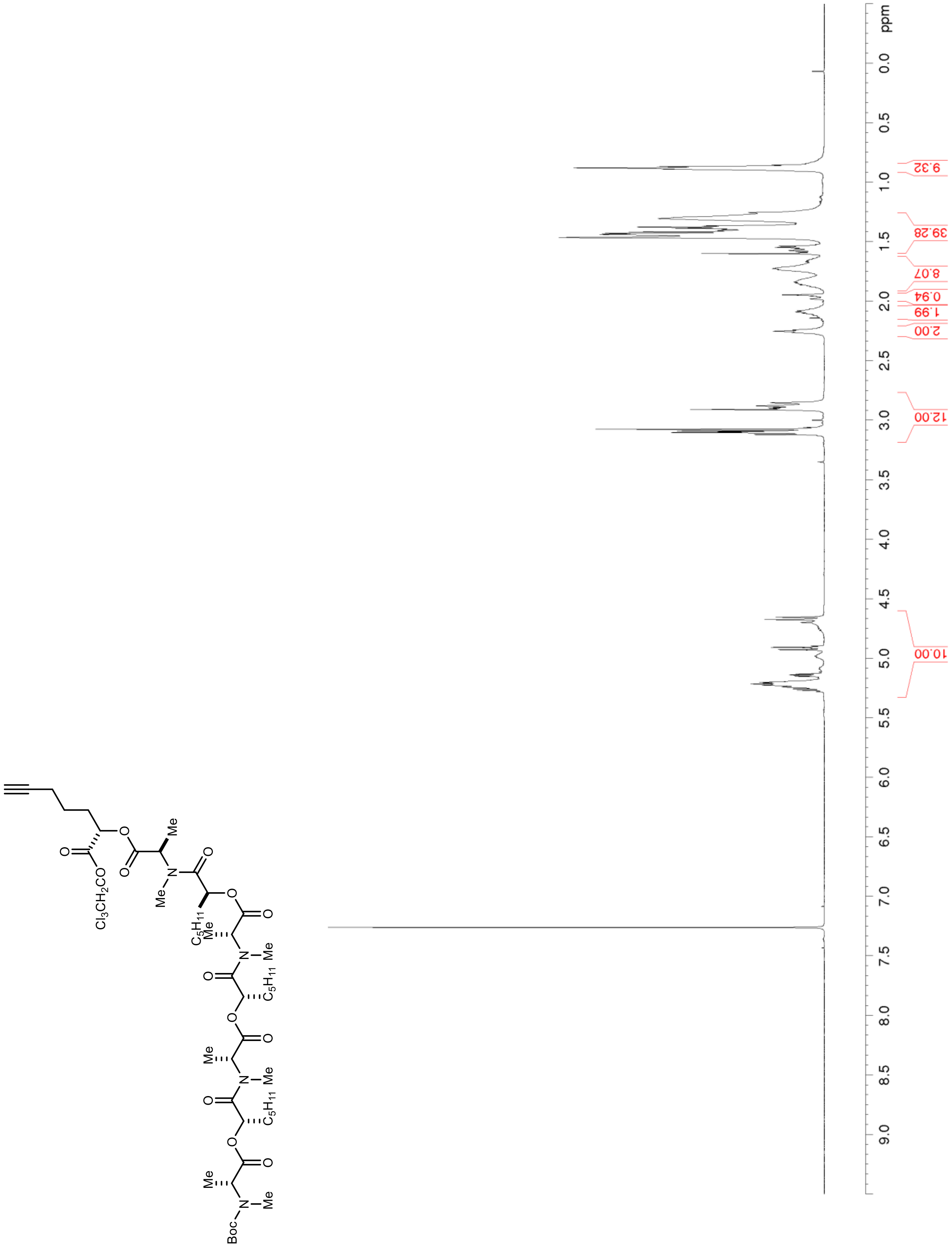

**Figure S13. $^{13}C$ NMR/DEPT (150 MHz, $CDCl_3$) of 12**

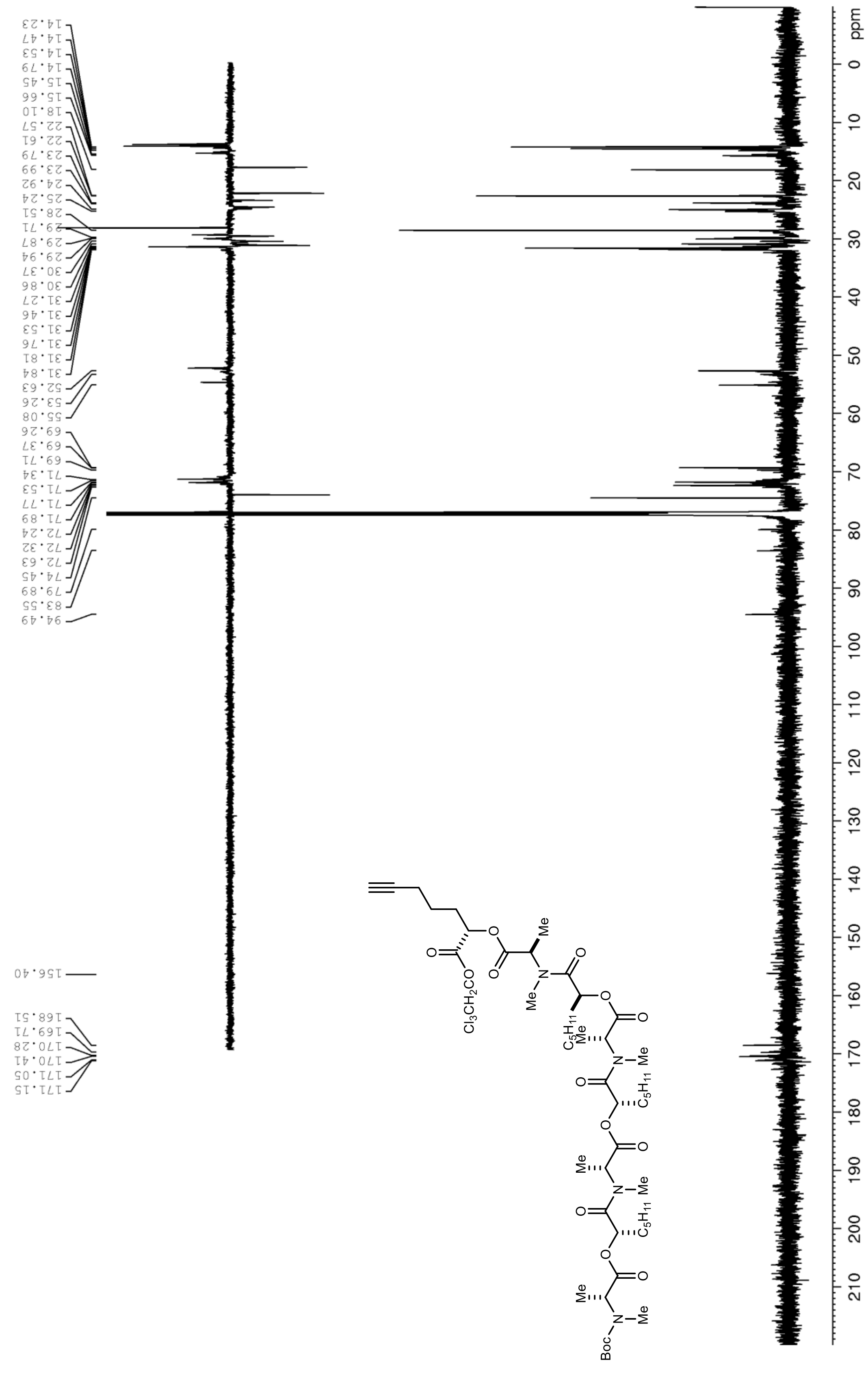

**Figure S14. $^1$H NMR (400 MHz, $CDCl_3$) of S2**

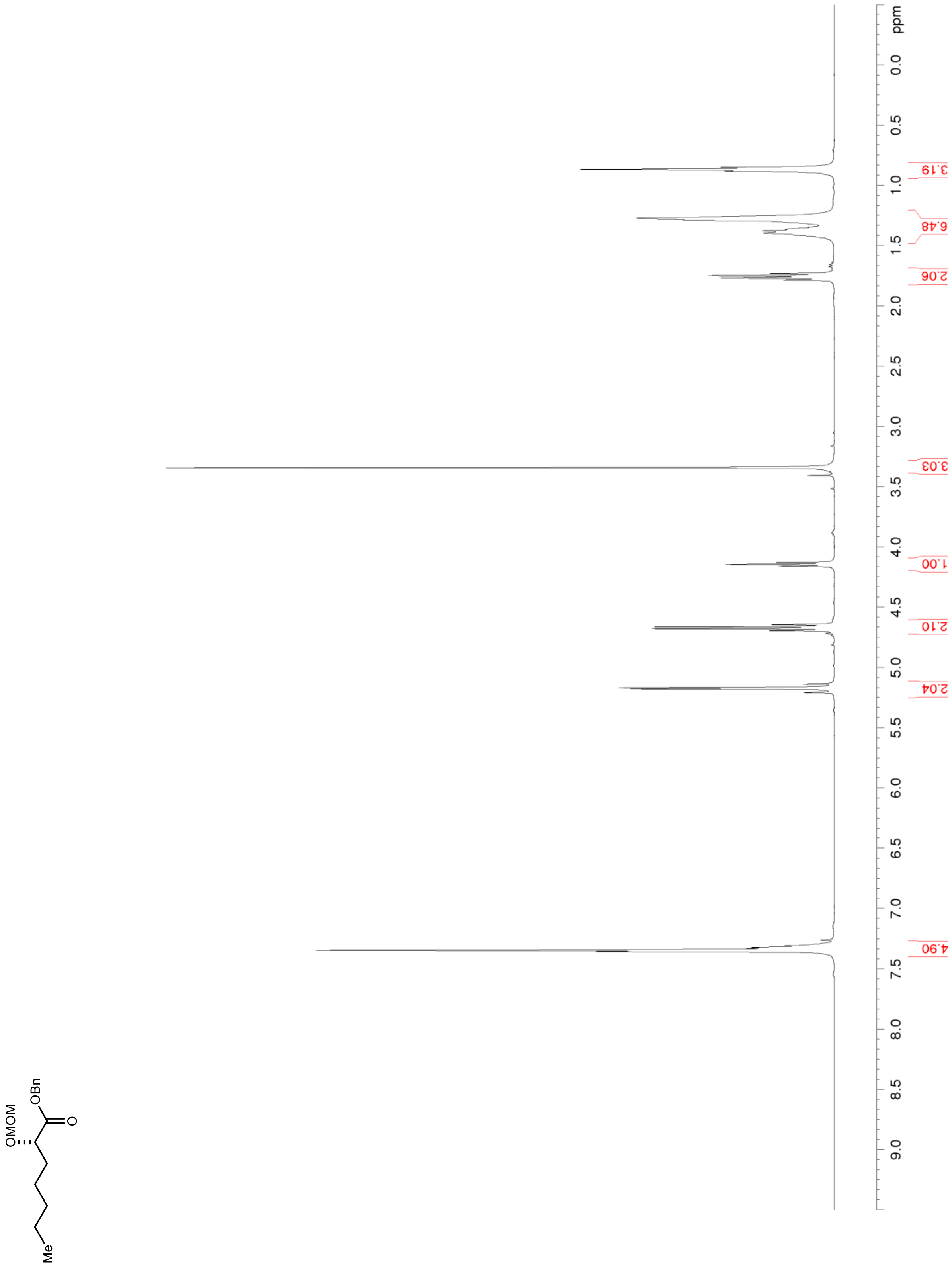

**Figure S15. $^{13}C$ NMR/DEPT (100 MHz, $CDCl_3$) of S2**

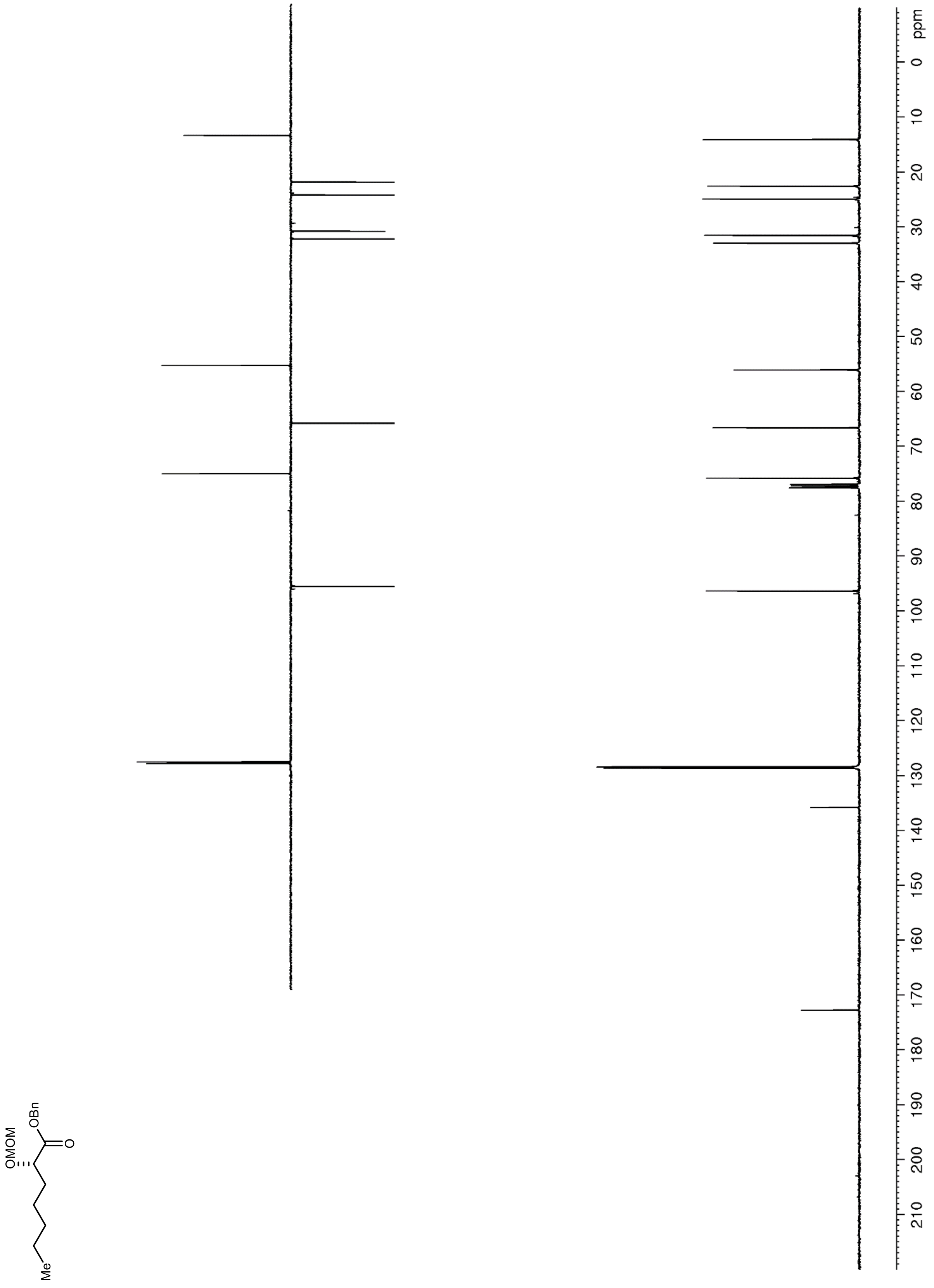

**Figure S16. $^1$H NMR (600 MHz, $CDCl_3$) of S5**

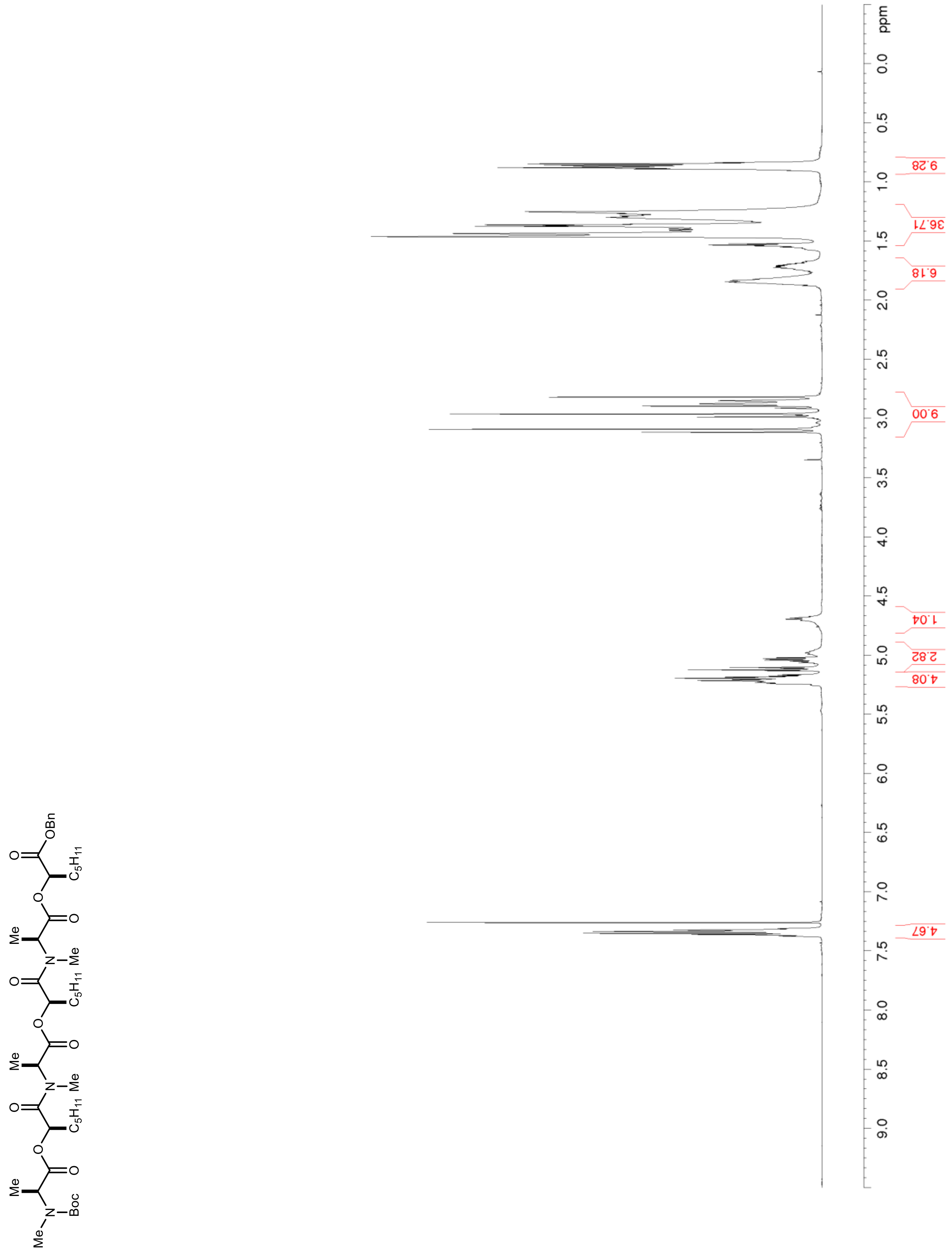

**Figure S17. $^{13}$C NMR/DEPT (150 MHz, $CDCl_3$) of S5**

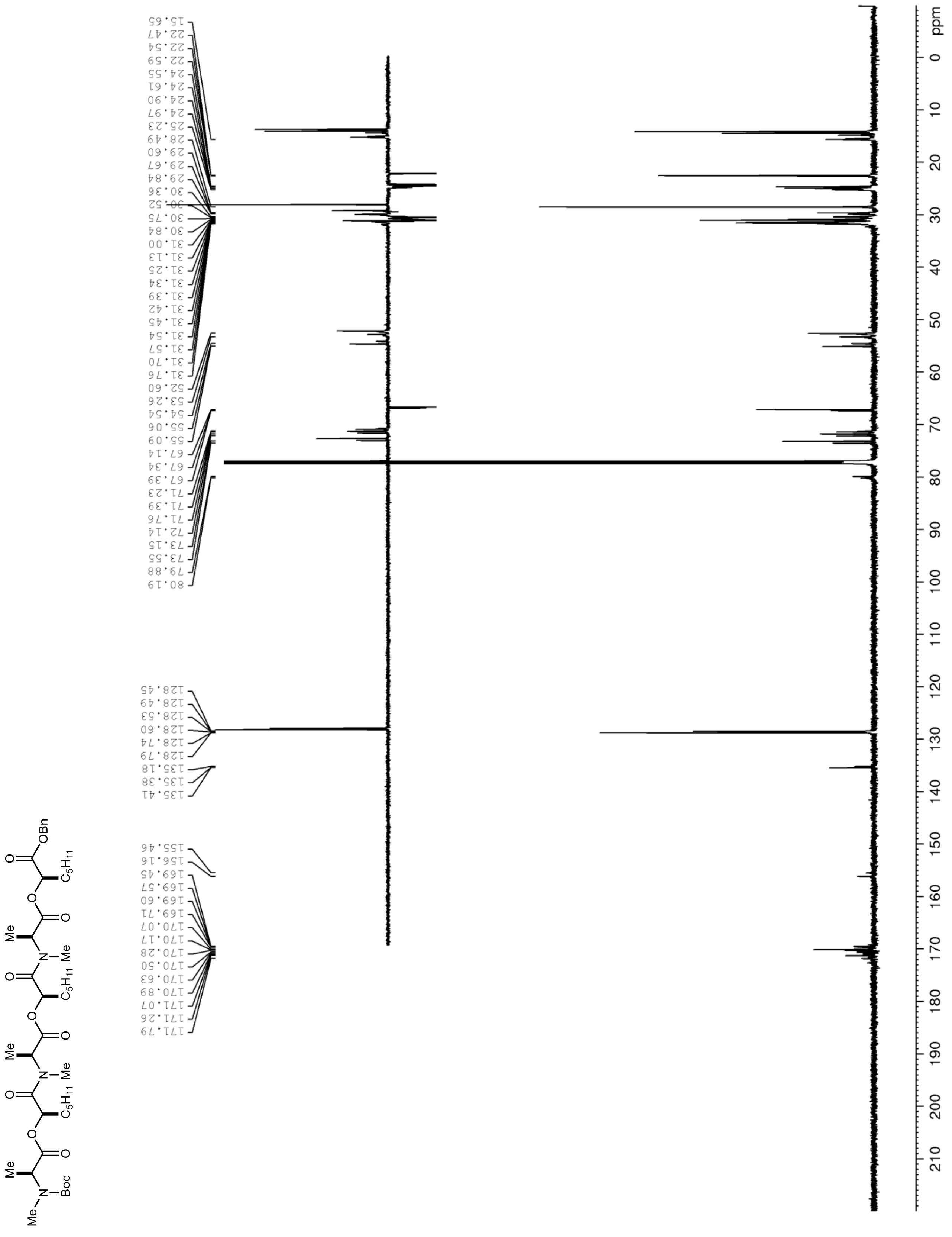
